# A phage parasite deploys a nicking nuclease effector to inhibit replication of its viral host

**DOI:** 10.1101/2021.07.12.452122

**Authors:** Kristen N. LeGault, Zachary K. Barth, Peter DePaola, Kimberley D. Seed

**Affiliations:** Department of Plant and Microbial Biology, University of California, Berkeley, 271 Koshland Hall, Berkeley, CA 94720, USA; Chan Zuckerberg Biohub, San Francisco, CA 94158, USA

## Abstract

PLEs are phage parasites integrated into the chromosome of epidemic *Vibrio cholerae*. In response to infection by its viral host ICP1, PLE excises, replicates and hijacks ICP1 structural components for transduction. Through an unknown mechanism PLE prevents ICP1 from transitioning to rolling circle replication (RCR), a prerequisite for efficient packaging of the viral genome. Here, we characterize a PLE-encoded nuclease, NixI, that blocks phage development likely by nicking ICP1’s genome as it transitions to RCR. NixI-dependent cleavage sites appear in ICP1’s genome during infection of PLE(+) *V. cholerae*. Purified NixI demonstrates *in vitro* specificity for sites in ICP1’s genome and NixI activity is enhanced by a putative specificity determinant co-expressed with NixI during phage infection. Importantly, NixI is sufficient to limit ICP1 genome replication and eliminate progeny production. We identify distant NixI homologs in an expanded family of putative phage satellites in Vibrios that lack nucleotide homology to PLEs but nonetheless share genomic synteny with PLEs. More generally, our results reveal a previously unknown mechanism deployed by phage parasites to limit packaging of their viral hosts’ genome and highlight the prominent role of nuclease effectors as weapons in the arms race between antagonizing genomes.

## INTRODUCTION

Biological conflicts represent a major selective pressure shaping microbial diversity from the level of genes to communities (1, 2). Nowhere are these conflicts more antagonistic than between bacteria and the lytic viruses (phages) that infect them. Bacterial genomes contain abundant anti-phage defense systems that function through diverse mechanisms to restrict phages. In addition to the selective pressure imposed by an evolving repertoire of host immunity, phages themselves are hosts to their own subcellular parasites. Phage parasites, often referred to as phage satellites, may appear to be defunct prophages, as they are integrated into specific attachment sites in their bacterial hosts’ genomes (3), yet phage parasites conspicuously lack the full suite of structural genes required for virion assembly (4). In response to proteins encoded by their viral hosts, phage parasites excise (5), replicate (6) and steal structural components from their viral hosts to package their own genome, and upon lysis of the cell, disseminate their genome to naïve neighboring cells in modified virions (7, 8). To accomplish this, phage parasites encode mechanisms to inhibit packaging of their viral host’s genome and favor packaging of the parasite genome. These mechanisms must balance inhibition of their viral host’s lifecycle with the phage parasites’ requirement for products encoded by the host virus. In this way, phage parasites employ finely tuned inhibitory proteins to redirect viral host resources for the parasites own use. Much of our understanding of the mechanisms that phage parasites use to subvert their viral hosts comes from the well-characterized phage-inducible chromosomal islands (PICIs) (4, 9), of which, <u>*S*</u>*taphylococcus* <u>*a*</u>*ureus* <u>p</u>athogenicity <u>i</u>slands (SaPIs) are the archetypical members (10). However, diverse families of unrelated phage parasites have likely evolved independently (11) and though often overlooked, such parasites are likely ubiquitous within bacterial genomes (12). Understanding diverse phage satellites can help define mechanistic paradigms of phage parasitism and uncover novel innovations not seen in model systems interrogated thus far.

PLEs (<u>p</u>hage-inducible chromosomal island-<u>l</u>ike <u>e</u>lements) are phage parasites unrelated to SaPIs that have proven lucrative for uncovering new mechanisms of parasitism (13–15). Based on current data, PLEs are restricted to toxigenic *Vibrio cholerae*, the causative agent of the diarrheal disease cholera. PLEs appear to exclusively parasitize the lytic phage ICP1, which is also frequently shed in cholera-patient stool samples (16). PLEs have been identified in *V. cholerae* genomes dating back to the 1940s, and to date five distinct PLEs have been identified that share considerable nucleotide identity (17). All PLEs are ~17kB in size and encode ~26 open reading frames (ORFs) that largely lack predicted functions or known PFAM domains. Recent surveillance efforts indicate that PLE 1 is currently the dominant PLE in *V. cholerae* isolated from cholera patients in Bangladesh where cholera is endemic (14, 17). For this reason, PLE 1 has been the subject of mechanistic studies aimed at understanding PLE activity (13, 15, 18, 19) and will generally be referred to here as ‘PLE’ unless another variant is specified. Although PLE is known to completely abolish ICP1 production (17, 20), the mechanisms PLE uses to inhibit ICP1 remain somewhat elusive. PLE employs multiple strategies to inhibit ICP1 progeny production, as no single ORF knockout allows ICP1 to overcome PLE (15). To antagonize PLE in nature, ICP1 isolates encode anti-PLE effectors: either a CRIPSR-Cas system that targets the PLE genome (20) or a site-specific nuclease directed to the PLE origin of replication (21). Such phage-encoded anti-PLE nucleases speak to the fitness costs ICP1 faces when PLE activity is left unimpeded.

Like other known phage parasites such as SaPIs, PLE modifies its viral host’s capsid, constraining ICP1’s capsid to ~1/3 of its normal size to accommodate the smaller PLE genome and exclude ICP1’s larger ~125 kB genome (13). Beyond redirecting virion morphogenesis proteins to package its own genome and restrict ICP1, recent work shows that PLE uses a multi-pronged approach to attack and compete with ICP1 DNA replication, both by reducing total ICP1 genome copy and by inhibiting a key step in ICP1’s genome replication program (18). PLE replication requires the PLE-encoded replication initiation factor RepA and hijacking of components of ICP1’s replisome to drive a nearly ~1000-fold increase in genome copy during ICP1 infection (17, 22). Concurrent with robust levels of PLE replication, PLE reduces ICP1’s DNA replication ~four-fold, suggesting a link between PLE replication and inhibition of ICP1 replication (18). In addition to restricting ICP1’s genome replication, PLE also alters the mode of ICP1 replication (18). At 8 minutes post-infection (mpi) of a permissive PLE(−) host, ICP1 replication proceeds through bidirectional theta replication (18). Later in infection, between 12 and 16 mpi, ICP1 transitions to rolling circle replication (RCR) (18). RCR is a common mode of replication for viral genomes and plasmids (23–25). Continued rounds of RCR generates linear genome concatemers, an essential step for many viruses because linear concatemers are the DNA substrate recognized by the terminase complex that initiates DNA packaging into capsids (26). When ICP1 infects a PLE(+) *V. cholerae* host, early bidirectional theta replication proceeds normally, however, ICP1’s transition to RCR is inhibited by an unknown mechanism (18). Importantly, even when PLE is unable to replicate (as in PLE (+)Δ*repA*) and total ICP1 replication is partially restored, ICP1’s transition to RCR is still inhibited, suggesting that an additional PLE-encoded factor impedes ICP1’s transition to RCR (18). To our knowledge, PLE is unique among phage parasites in targeting genome replication of its viral host, highlighting this stage of the viral lifecycle as a ripe arena for uncovering new mechanisms of interference.

Here, we work to uncover the PLE-encoded mechanism that inhibits ICP1’s RCR. We identify aberrant cut sites in ICP1’s genome that appear during infection of a PLE(+) host and identify the PLE-encoded nuclease necessary for generating these cut sites. This PLE-encoded nuclease is co-expressed with a small protein coincident with ICP1’s attempt to transition to RCR in a PLE(+) host. We provide evidence that this small protein acts as a specificity factor, increasing the nuclease’s activity and binding to regions of ICP1’s genome shown to be cut *in vivo*. We further identify homologs of the PLE-encoded nuclease in what we consider to be an expanded family of putative phage parasites in non-toxigenic *V. cholerae* and other non-cholera Vibrio species.

## MATERIALS AND METHODS

### Strains and media

A complete list of strains used in this study is available in Supplementary Table S1. *V. cholerae* strains used in this study are derivatives of E7946. ICP1_2006_E engineered to lack CRISPR-Cas (ΔCRISPR, Δcas2-3) was used throughout this study and is referred to ICP1 for simplicity (19). Routine culturing of *V. cholerae* was done in LB at 37°C with aeration. Cells grown in Terrific Broth (TB) were used for protein purification. Antibiotics were supplemented as appropriate at the following concentrations: 50 μg/ml kanamycin for maintenance of the pSUMO vector, 100 μg/ml spectinomycin, 1.25 μg/ml or 2.5 μg/ml chloramphenicol for broth or plate conditions, respectively.

### Cloning

Plasmids to express *nixI* and *stiX* are pMMB67EH derivative plasmids with a theophylline inducible riboswitch (Riboswitch E) as described previously (19), and were constructed using Gibson assembly. For expression of *nixI* homologs from *V. cholerae* YBA_A06 (WP_057552372) and *Vibrio parahaemolyticus* S042 (WP_029837635), the genes were commercially synthesized by Genscript and cloned into the pMMB67EH derivative. High copy pUC19 derivative plasmids were used as DNA substrates for nuclease assays. Deletions in *V. cholerae* were generated through natural transformation as described previously (27). Naturally competent *V. cholerae* were transformed with PCR products generated by splicing by overlap extension PCR containing arms of homology flanking frt recombination sites as described previously (27). All deletions and constructs were verified by Sanger sequencing.

### Protein purification

Protein purifications were carried out in Buffer A (150mM Tris-Cl pH=7.8, 50mM NaCl, 2mM BME, 10mM NaCl) unless otherwise indicated or modified. His-SUMO tagged NixI^N95A^ and StiX were purified from *Escherichia coli* strain BL21. Catalytically active His-SUMO tagged NixI was purified from *E. coli* BL21 pLysS background to reduce leaky expression of *nixI* which is toxic to *E. coli*. For each *E. coli* expression strain, a 100mL starter culture in TB was grown overnight at 37°C on a shaking incubator at 250 rpm with the appropriate antibiotic(s): kanamycin at 50 μg/ml to maintain the pSUMO plasmid, and chloramphenicol at 25 μg/ml to maintain the pLysS plasmid. The next day, four 1L baffled flasks of TB with appropriate antibiotic(s) were each inoculated with 20mL of overnight starter culture, and grown at 37°C with shaking. When the OD_600_ reached 0.5-0.7, the temperature of the incubator was reduced to 18°C, 0.5mM IPTG was added to induce protein expression, and cultures were incubated for 20 hours with shaking. Cells were harvested by centrifugation at 4000xg for 15 minutes then resuspended in 25mL of lysis buffer: 150mM Tris-HCl pH=8, 50mM NaCl, 2mM BME, 1X Pierce phenylmethylsulfonyl fluoride protease inhibitors + 0.5% Triton-X, 1% glycerol. The cell suspension was sonicated for a total of 5 minutes as cycles of 10-second “ON” pulses with 20-seconds “OFF”. After sonication, cell debris was removed by centrifugation at 20,000xg for 40 minutes, the supernatant was filtered using GE 0.2μm regenerated cellulose membranes. Lysate was bound to a GE His-Trap and pumped through using a peristaltic pump. A high salt wash including 2M NaCl was applied to remove DNA. The His-SUMO tagged proteins were eluted in 250mM imidazole in 150mM Tris-Cl pH=7.8, 50mM NaCl, 2mM BME. The eluted protein was then dialyzed against Buffer A + 5% glycerol overnight using SnakeSkin (Thermo Scientific) at 10kDa MWCO dialysis membrane for NixI and NixI^N95A^ or 3kDa MWCO dialysis membrane for StiX. Dialysis was performed in concert with cleavage of the His-SUMO tag using the SUMO protease (Sigma Aldrich) at 20U/1mg of protein. To remove the SUMO protease and the cleaved tag, the protein was re-applied to Ni-NTA resin and a batch purification was performed, where the cleaved protein of interest remained unbound in the buffer. Protein concentration was measured using a BioPhotometer D30 (Eppendorf). Additional purification of the Ni-affinity purified NixI and NixI^N95A^ was carried out using a Heparin column, equilibrated with Buffer A except with a higher NaCl concentration of 150mM. The column was washed with Buffer A+ 150mM NaCl until no protein was detected in the flow through, then eluted with 2M NaCl in Buffer A, and dialyzed against Buffer A overnight. Additional purification of StiX was performed using size exclusion chromatography on the GE AKTA, with fractions containing StiX pooled and concentrated using a 3kDa MWCO centrifugal filter.

### Electrophoretic mobility shift assays (EMSAs)

NixI^N95A^ with and without StiX were incubated with probes amplified from *gp121* or a control region in ICP1 from *gp62*. Primer sequences are available in Supplementary Table S2. 20nM of DNA probe was added for each 20μL reaction, and incubated with varying concentrations of NixI and StiX, either alone or in combination. Reactions were carried out for 30 minutes at 30°C and then chilled at 4°C. Reactions were run on an 8% polyacrylamide gel, pre-run at 120V for 30 minutes. EMSA reactions were run at 120V for 30 minutes at 4°C, then visualized by staining with GelRed.

### Nuclease assays

As HNH nucleases require a divalent metal co-factor for activity (typically magnesium, manganese or zinc (28–31)), we investigated NixI’s metal requirement with a panel of divalent metals at differing concentrations and determined NixI can utilize Mg^2+^ or Mn^2+^ for catalysis, but not Zn^2+^ or Ca^2+^ (Supplementary Figure S1). Although Mn^2+^ can stimulate nuclease activity of NixI, work with other HNH nucleases, including a sequence specific nicking endonuclease (32), has shown that Mn^2+^ can replace Mg^2+^, leading to non-specific activity (33, 34). Therefore, we carried out our nuclease reactions in a buffer supplemented with 10mM Mg^2+^. Nuclease assays with NixI and NixI^N95A^ were carried out in 150mM Tris-Cl pH=7.8, 50mM NaCl, 1mM DTT and 10mM MgCl_2_. Reactions containing StiX were performed with a constant amount of StiX at 500nM. Plasmid DNA substrates were added at 100ng per 20μl reaction, with NixI at varying concentrations as indicated. Reactions were incubated at 30°C for 15 minutes, then 1μg/mL of Proteinase K was added to each reaction to digest bound protein and incubated at 37°C for 30 minutes. The entire reaction volume was then run on a 0.8% agarose gel stained with GelRed. To generate cleaved controls, 100ng of vector was digested for 1 hour at 37°C with 10 U of BamHI-HF (NEB) to create a single double-stranded cut or 10 U Nb.BtsI (NEB) to create a single nick, then the enzymes heat inactivated at 80°C for 20 minutes.

Densitometry measurements were performed using ImageJ to quantify nuclease activity on plasmid substrates. Percent cleaved was calculated as the intensity of the supercoiled+linearized plasmid divided by the total band intensity in each lane. Densitometry measurements reported here are the average of three replicate nuclease assays conducted on different days.

### Sample preparation for DNA sequencing and reads mapping

ICP1 infected PLE(+)Δ*nixI V. cholerae* samples were prepared in parallel with previously analyzed PLE(+) and PLE(−) *V. cholerae* samples, as described previously (18). Briefly, 2ml LB cultures of each *V. cholerae* strain was infected at OD_600_=0.3 with ICP1 at a multiplicity of infection (MOI) of 1, and aliquots taken at given time points from which total DNA was extracted for sequencing. The pellets were washed with 1mL ice cold phosphate buffered saline, and total DNA was extracted using the QIAGEN DNeasy Blood and Tissue kit. Library prep was done using the NEBNext Ultra II FS DNA Library Prep Kit, and sequenced using Illumina HiSeq4000 at University of California Berkeley QB3 Core Facility. Illumina sequencing reads for each sample and time point were mapped as before (18). Briefly, reads were mapped onto ICP1 reference genome using Bowtie 2 v2.3.4.1 using the default settings except for “-end-to-end’ and “-very-sensitive”. Read coverage was normalized by the total number of reads that mapped to the reference.

### *In vivo* cleavage site identification

Illumina reads from samples at 16 minutes post-infection were analyzed for indications of DNA cleavage from PLE(−), PLE(+) and PLE(+)Δ<u>*nixI*</u> *V. cholerae*. Cleavage ratios in ICP1’s genome were calculated following an approach adapted from Culviner et al. (35). To look for putative cleavage sites, the log_2_ ratio of reads coverage averaged across three biological replicates at each nucleotide position in ICP1 was calculated for the following comparisons: PLE(+):PLE(−) and PLE(+)Δ*nixI*:PLE(−). The difference in log_2_ coverage was determined for each nucleotide position by subtracting the coverage at the nucleotide before and after each position to determine coverage decrease in both a 5’ to 3’ and 3’ to 5’ orientation. These differences were then ranked, identifying 63 putative sites with coverage change >log_2_(0.026) (Supplemental Figure S2).

### Efficiency of center of infection assays

Cultures of *V. cholerae* were grown in LB supplemented with chloramphenicol (Cm) to maintain the plasmid expressing NixI, NixI+StiX or the empty vector control. At an OD_600_=0.2, expression from the plasmid was induced with 1mM IPTG + 1.5mM Theophylline. At OD_600_=0.3, approximately 15 minutes post-induction, 2mL cultures were infected at MOI=0.1 and incubated for 7 minutes to allow adsorption of ICP1. The infected culture was then diluted 1:2500, 1:25000 and 1:250000 into pre-warmed LB supplemented with Cm and inducer to a final volume of 2ml. Immediately after dilution, 500μl of 1:2500 diluted sample was removed to quantify the input. To quantify phage produced, 500μl of each dilution was collected 40 minutes after the initial infection. To all samples, 20μl of chloroform was added to each sample and debris and chloroform was removed by centrifugation at 5,000xg for 15 minutes. Supernatant was used to quantify phage by plaque assay on a permissive PLE(−) host. Efficiency of center of infection was quantified as the ratio between the number of phages produced ml^−1^minus the number of input phage, from the strains of interest compared to the control culture: either an empty vector control when assessing restriction by *nixI*+*stiX* or wild type PLE.

### Quantification of genome replication by real-time quantitative PCR

ICP1 and PLE replication was quantified by qPCR as described previously, where strains used for quantifying ICP1 replication were infected at an MOI=0.1 and strains used for quantifying PLE replication infected at an MOI=2.5 (17). For strains expressing plasmid constructs, a 2mL culture of interest grown with chloramphenicol was induced with 1mM IPTG 1.5mM theophylline at OD_600_=0.2, approximately 15 minutes before infection when OD_600_=0.3. A T=0 aliquot was immediately boiled to stop DNA replication. Cultures were incubated for 20 minutes at 37°C with aeration, then a 20μl aliquot was boiled. DNA replication was determined by quantifying the fold change in replication at T=20 compared to genome copy at T=0. ICP1’s genome was amplified with primers Zac68/69 and PLE replication with primers Zac14/15 using iQ SYBR Green Supermix (Bio-Rad) and the CFX Connect Real-Time PCR Detection system (Bio-Rad). Replication efficiency was compared to a PLE(−) permissive strain harboring an empty vector and similarly grown in the presence of chloramphenicol and inducer.

### Identification of an expanded family of PLEs

An HHBlits search with PLE 1’s NixI protein sequence returned significant homology across the protein sequence to a protein found in non-O1 *V. cholerae* (WP_057552372). The identical protein was found in a total of 10 non-O1 *V. cholerae* isolates isolated in Massachusetts. Analysis of the 10kb regions flanking the NixI homolog showed shared synteny with PLEs, specifically these regions are characterized by a 5’ integrase, a total sequence length of ~17kB, and an ORF-less region between divergently transcribed operons. To identify additional PLEs, a PSI-BLAST search excluding *V. cholerae* was performed with NixI from *V. cholerae*’s PLE 1 as well as the NixI homolog from non-toxigenic *V. cholerae* YB2_A06 (WP_057552367.1). This revealed a family of homologs in other Vibrios, such as *V. alginolyticus*, *V. parahaemolyticus*, and *V. casei*, sharing around 30% amino acid identity across >85% of the protein length. These Vibrio NixI homologs were found in a similar syntenic context as NixI from PLE, but with a 5’ upstream tyrosine recombinase in place of the large serine recombinase characteristic of *V. cholerae* PLEs. All PLEs had a predicted replication initiation protein in the same syntenic location as PLE 1’s RepA: either a RepA_N homolog or DNA Pol I C-terminal domain-containing protein. Notably, all homologs of NixI found outside of toxigenic *V. cholerae* PLEs lacked a small downstream ORF with homology to StiX. To ensure that these annotations were called correctly, the PLEs from *V. parahaemolyticus* S042 and *V. diabolicus* were verified with RAST. Additionally, the start codon for the downstream peptidase was only a few nucleotides downstream of the end of the NixI homolog, further supporting the lack of a StiX homolog in PLEs outside of *V. cholerae*.

### Statistical and visual analyses

Statistics and graphs were generated using Prism version 9.0.2. Gene maps were made using the gggenes package in R, version 4.0.0. Consensus motifs of NixI cleavage sites were produced by the WebLogo server (https://weblogo.berkeley.edu/logo.cgi).

## RESULTS

### ICP1’s genome is cleaved during infection of PLE(+) *V. cholerae*

While assessing ICP1 replication profiles by mapping Illumina read coverage for each nucleotide position (referred to from here on out as DNA-seq) (18), we noted regions in ICP1’s genome where reads coverage dropped precipitously during infection of PLE (+) but not PLE(−) *V. cholerae*. These coverage drops suggested potential sites of nuclease cleavage. In order to characterize these putative cut sites more systematically we drew on a previously published method that identified cleaved sequences in transcripts from RNA-seq data in order to assess activity of an endoribonuclease toxin (35). To generate cleavage maps across ICP1’s genome, we calculated the coverage ratio of ICP1 reads 16 minutes post-infection of PLE(+) relative to PLE(−) *V. cholerae*. This time point is late in the infection cycle, at a time when ICP1 and PLE are both replicating (18) and before the onset of lysis ~20 minutes post-infection of PLE(+) *V. cholerae* (15, 17) (Figure 1A). We identified a list of PLE-dependent candidate cut sites in ICP1 with log_2_ coverage drops >(0.026) for a total of 63 sites, with several coverage drops clustering in the same region of ICP1’s genome (Materials and Methods, Supplemental Figure S2). Two sites in ICP1’s genome, located in *gp121* and *gp156*, showed the most striking drops in coverage, while additional sites showed more subtle drops in coverage (Figure 1B and 1C, Supplemental Figure S2). We therefore chose to focus on the sites in *gp121* and *gp156* in greater detail. We identified a shared motif of GTAAWC^TT, where the “^” denotes the position of potential cleavage (Figure 1D). Where multiple coverage drops occurred within a 100bp region (as was common for a few regions, Supplemental Figure S2), the sequence at the most dominant coverage drop was used as input to generate the consensus motif for all 63 sites. These sites generated a less stringent motif GNAANC^TW that occurs 99 times in ICP1, of which 16 sites displayed a coverage drop at the 16 minute time point (Supplemental Figure S2). The sites in ICP1’s genome that have the motif but were not cleaved may not be cleaved due to protein occupancy or were perhaps missed because this assay was performed at just one time point, though additional DNA features such as proximal sequence or specific DNA topology may also contribute to cleavage. The analysis of reads coverage across ICP1’s genome suggests a limited number of pronounced coverage drops that appear only when infecting a PLE(+) host, consistent with nucleolytic cleavage at these regions.

**Figure 1.**
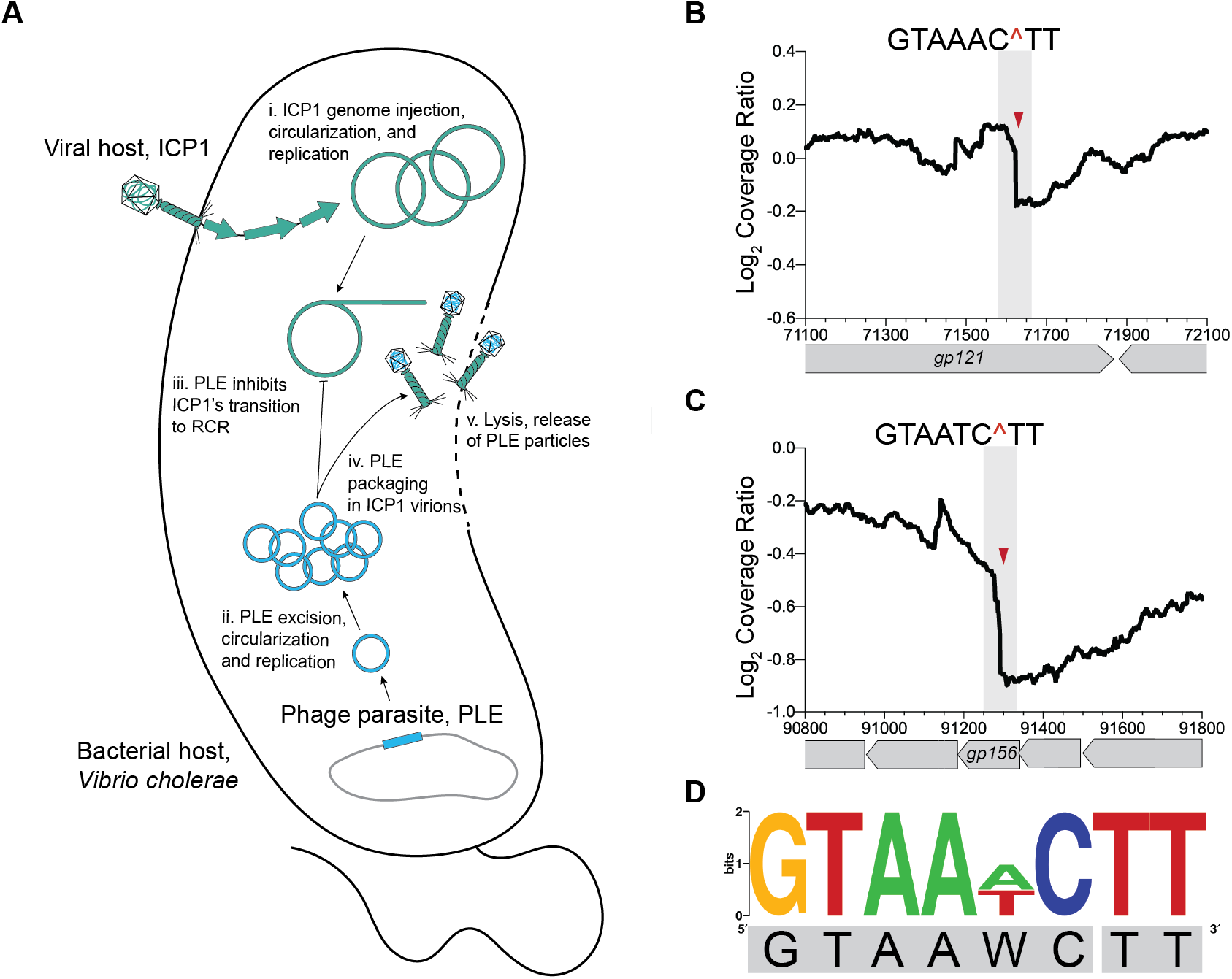
ICP1’s genome exhibits signs of cleavage when infecting PLE(+) *V. cholerae*. (A) Model of ICP1 infection of a PLE(+) *V. cholerae*: The viral host ICP1 injects its genome, triggering PLE excision and gene expression. Both ICP1 and PLE replicate their genomes initially, yet midway through infection PLE prevents a further increase in ICP1 genome copy and inhibits ICP1’s transition to rolling circle replication (RCR). PLE hijacks ICP1 structural proteins for packaging, and no ICP1 progeny are produced. *V. cholerae* lyses ~20 minutes post-infection releasing only PLE transducing particles. (B) Example cleavage ratio PLE(+)/PLE(−) across ICP1’s genome position (x-axis) showing the putative cut site within ICP1’s *gp121*, with the cut position indicated by the red triangle and the nucleotides flanking the cut region above. The cleavage ratio is calculated as the average coverage of three biological replicates. (C) As in B, but displaying the next most prominent cut site within ICP1’s *gp156*. Additional cut sites are summarized in Supplemental Figure S2. (D) Weblogo displaying the consensus motif at cut sites in *gp121* and *gp156*.

### ICP1’s genome is targeted by a PLE-encoded nicking endonuclease, NixI

We hypothesized that the putative cleavage sites in ICP1 apparent during infection of PLE(+) *V. cholerae* were caused by a PLE-encoded nuclease. However, we lacked a candidate nuclease, as the majority of PLE-encoded open reading frames (ORFs) have no primary homology to any annotated genes. Using the more sensitive HHpred, which searches for remote homology based on predicted structural domains, we identified a single PLE-encoded ORF, NixI (WP_032468646), that shares homology with the intron-encoded homing endonuclease I-HmuI (probability score=97.1%, e-value=0.0058) (30). Despite sharing only 10% amino acid identity across the protein, NixI shares an HNH/N active site domain with I-HmuI (Figure 2A). No other proteins shared significant homology with NixI by HHpred, which only compares proteins with solved structures. I-HmuI is the only crystallized protein from a family of closely related nicking HNH/N endonucleases (30, 36), and a MUSCLE alignment including these proteins identified several conserved residues at the catalytic core, including an experimentally validated catalytic asparagine from I-HmuI at position 95 in NixI (30) (Figure 2A). To address whether *nixI* may encode for a nuclease, we cloned *nixI* or a mutated *nixI*^N95A^ allele, which we postulated would be catalytically inactive, under an inducible promoter in *V. cholerae* and plated for cell viability with and without inducer. Consistent with *nixI* functioning as a nuclease, induction completely abolished *V. cholerae* colony formation, as the predicted recognition motif we identified in ICP1’s genome, (GTAAWC^TT), is found 246 times in *V. cholerae*’s chromosomes. In contrast, induction of NixI^N95A^ had no impact on colony formation (Figure 2B), indicating that toxicity of NixI is associated with nucleolytic activity.

**Figure 2.**
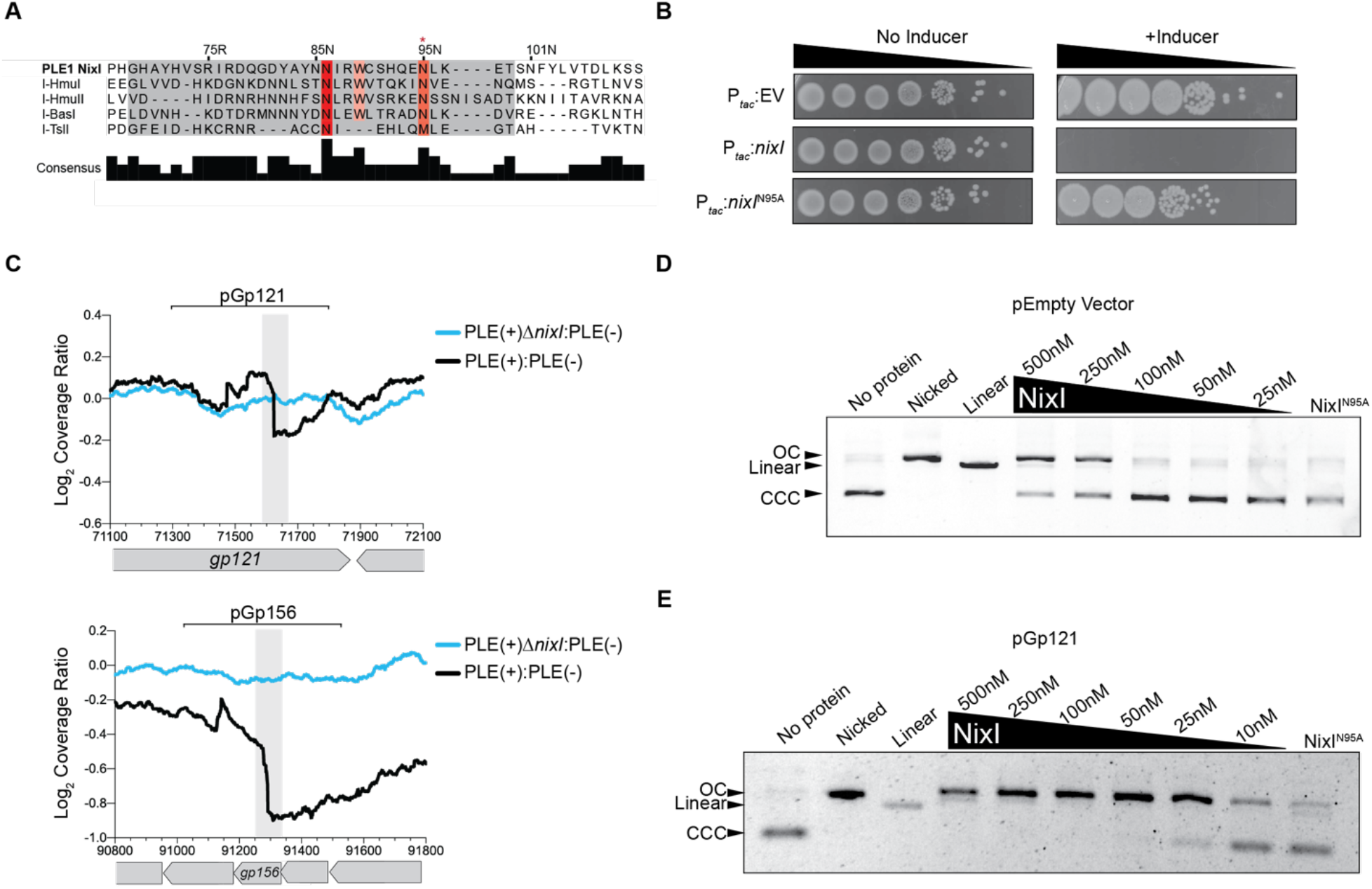
PLE encodes a nuclease, NixI, that cuts ICP1’s genome *in vivo* and *in vitro*. (A) MUSCLE alignment of the catalytic core of phage-encoded intron homing endonucleases including I-HmuI, where I-HmuI’s active site domain is highlighted in grey and residues that share >70% identity in orange/red. The red asterisk above 95N indicates the experimentally validated catalytic asparagine residue from I-HmuI (30). (B) 10-fold dilution series of mid-log grown *V. cholerae* containing plasmids for inducible expression of *nixI*, *nixI* with the putative catalytic asparagine mutated to an alanine (N95A), or an empty vector control, spotted on plates +/− inducer. (C) Coverage ratios PLE(+)/PLE(−) and PLE(+)Δ*nixI* /PLE(−) across ICP1’s putative cleaved genes *gp121* and *gp156* as indicated in Figure 1B, as the average coverage of three biological replicates. Brackets above indicate the regions cloned into the plasmid and used as a DNA substrate for *in vitro* assays. (D-E) *In vitro* cutting by purified NixI at the concentrations indicated or 500nM NixI^N95A^ of an empty vector plasmid (D) or a plasmid with a 500 base pair region from *gp121* as indicated in panel C (E). CCC=covalently closed circular DNA, OC=open circular DNA.

Having identified a putative PLE-encoded nuclease, we next asked whether *nixI* is required for the appearance of the putative cleavage sites in ICP1’s genome during infection of PLE(+) *V. cholerae*. We infected PLE Δ*nixI V. cholerae* with ICP1 and again performed DNA-seq to evaluate changes in cleavage ratios of PLE(+)Δ*nixI* relative to *V. cholerae* PLE(−). Strikingly, we observed that all coverage drops in ICP1’s genome seen during a PLE(+) infection were abolished during infection of PLE(+)Δ*nixI* (Figure 2C and Supplemental Figure S2), indicating that *nixI* is necessary for cleavage of ICP1’s genome.

To directly evaluate NixI’s nuclease activity, we purified NixI and NixI^N95A^ (Supplemental Figure S3) to conduct *in vitro* nuclease assays. Many nucleases require specific DNA topology such as negative supercoiling for activity, so we first assayed whether NixI had nucleolytic activity on plasmid substrates, allowing for visualization of both nicking, as evidenced by a loss of supercoiled plasmid, and double stranded breaks. We observed nuclease activity on an empty vector plasmid at high protein concentrations (250nM and greater) after a 15-minute incubation period (Figure 2D). We postulated that NixI exhibits a preference for cleavage of sites in ICP1’s genome that we identified coverage drops in during infection of a PLE(+) host (Figure 2C). To test this, a 500 bp region flanking the putative cut site in *gp121* was cloned onto the same plasmid and used as a substrate for NixI cleavage assays. NixI nucleolytic activity was dramatically enhanced for a plasmid template containing the region from *gp121*, allowing us to observe cleavage with only 25nM NixI (Figure 2E). Importantly, NixI^N95A^ did not exhibit any nuclease activity, showing that despite low primary sequence homology, NixI does indeed share a catalytic motif with I-HmuI (Figure 2A). We also assayed cleavage of the predicted cleavage site within *gp156* and observed similar levels of cleavage as we observed for *gp121* (Supplementary Figure S4A). To determine whether NixI is a nickase or double-stranded cutter, we compared cut products to plasmid controls that had been nicked or linearized using commercially available endonucleases with known sequence specificity. NixI generated one or more nicks in the DNA substrate but did not cause double stranded cuts (Figure 2E). We next assayed whether NixI cleaves PCR products. Even at high protein concentrations we observed little nucleolytic activity of PCR products, as visualized on native or denaturing agarose gels (Supplementary Figure S5). Together, these data show that PLE encodes a nicking endonuclease, NixI, that is necessary for cleavage of ICP1’s genome *in vivo* and that shows *in vitro* specificity for sequences found in ICP1’s genome. We propose that NixI is a PLE-encoded nuclease effector that targets ICP1, functioning as part of PLEs strategy to antagonize its viral host.

### NixI nuclease activity is modulated by a small co-expressed protein StiX

Previous RNA-seq analyses of ICP1 infecting PLE(+) *V. cholerae* indicated that PLE is transcriptionally activated upon infection by ICP1, though most PLE-encoded ORFs were expressed at low levels in the absence of infection (37). An exception is PLE-encoded *nixI*, which is completely transcriptionally repressed in the absence of ICP1 infection, consistent with NixI’s nuclease activity and toxicity to *V. cholerae* (Figure 2B). By RNA-seq, *nixI* is co-transcribed with a small downstream ORF, which we have named *stiX*, with peak expression of *nixI+stiX* observed between 12- to 16-minutes post-infection by ICP1 (37) (Figure 3A). To determine whether StiX functions in concert with NixI, the genes were cloned under an inducible promoter retaining the native orientation of *nixI* closer to the promoter. Strikingly, co-induction of *nixI*+*stiX* reduced *nixI*’s toxicity and rescued colony formation of *V. cholerae* (Figure 3B). To quantify *stiX*-mediated rescue, strains expressing either *nixI*, *nixI*+*stiX* or an empty vector were induced, and cell viability 30 minutes post-induction was determined. We observed that cell viability was increased ~100-fold when *stiX* was co-expressed (Figure 3C), suggesting that StiX may restrict NixI nuclease activity.

**Figure 3.**
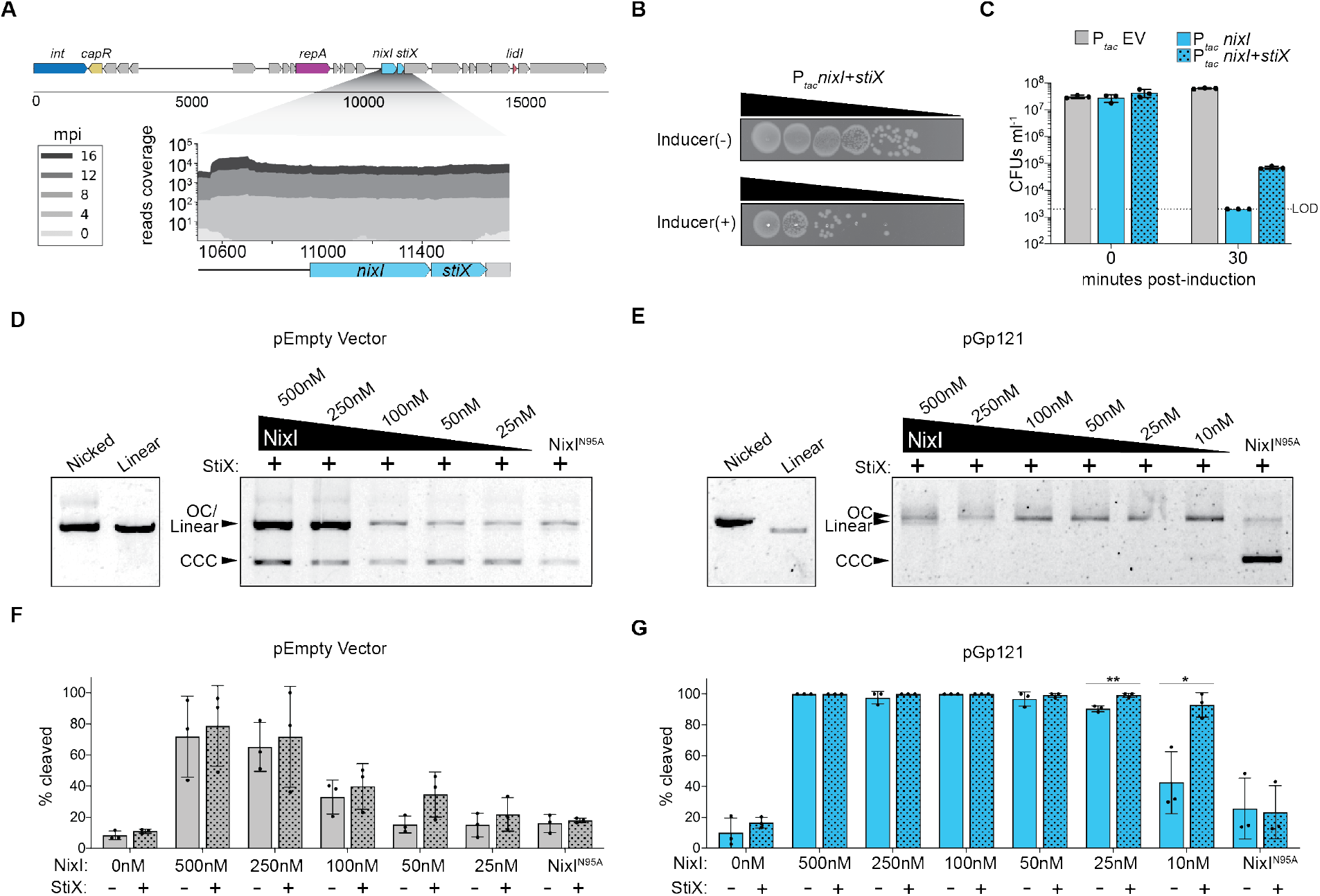
NixI’s nuclease activity is modulated by StiX. (A) PLE 1 gene map showing the relative position of *nixI* and *stiX* as well as characterized PLE genes, and previously published RNA-seq reads (37) of PLE 1 transcripts mapped onto this region showing co-transcription of *nixI* and *stiX* midway through infection by ICP1. (B) 10-fold dilution series of mid-log grown *V. cholerae* with inducible chromosomal constructs expressing *nixI*+*stiX* +/− inducer. (C) Cell viability as measured by CFUs ml^−1^ of *V. cholerae* with an empty vector (EV), or the genes indicated prior to induction and 30 minutes post-induction. Bar height displays the mean, whiskers are standard deviation, dots indicate independent biological replicates, LOD=limit of detection. (D-E) *In vitro* cutting by purified NixI/ NixI^N95A^ supplemented with 500nM StiX, showing cleavage of an empty vector (D) or plasmid containing *gp121*(E), with plasmid controls as in Figure 2D and E. (F-G) Densitometry measurements calculating the percent of cleaved plasmid out of total band intensity per lane. Plasmid cleavage of an empty vector (F) or plasmid with *gp121* (G) by NixI (solid bars) or NixI+StiX (patterned bars), or NixI^N95A^ (right most bars). Bar height is the mean, whiskers show standard deviation, dots are measurements from independent assays. Asterisks indicate statistical significance between NixI alone and NixI+StiX (*P<0.05; **P<0.01, where no values are shown no difference was detected, two-tailed t-test).

Like NixI, we were unable to detect any conserved domains nor homologs of StiX outside of PLEs using BLAST; however, using HHpred we found that StiX shares predicted structural homology with the C-terminal domain of *E. coli’s* alternative extracytoplasmic sigma factor SigE (Probability score=98.06%, e-value<0.0003, 21% total amino acid identity). The predicted structure of StiX generated by i-TASSER (38) shows homology between StiX and the σ^4^ domain of SigE, which contacts the −35 promoter element but does not participate in strand separation (Supplementary Figure S6) (39, 40). The high degree of predicted structural homology between StiX and SigE led us to hypothesize that StiX may alter NixI’s cutting by modulating its DNA binding. To test this, we purified StiX and assessed its impact on NixI-mediated cleavage of plasmid substrates. With 500nM StiX and a decreasing concentration of NixI, we again saw weak nicking of an empty vector plasmid (Figure 3D). Surprisingly, we saw that StiX increased NixI nuclease activity on a substrate possessing the region from ICP1’s *gp121* (Figure 3E). To assess the effect of StiX on NixI activity more quantitatively, we measured the amount of cleaved product relative to total band intensity in each lane of the gel for DNA substrates incubated with NixI, StiX, NixI+StiX or NixI^N95A^ (Figure 3F–G). StiX did not augment NixI cleavage of the empty vector template (Figure 3F). Additionally, StiX did not exhibit nuclease activity on its own (Figure 3G) nor did incubation with NixI^N95A^ result in cleavage (Figure 3F–G). However, StiX enhanced NixI’s nucleolytic activity on a substrate containing *gp121*, significantly increasing the amount of cut substrate at NixI concentrations under 50nM (Figure 3G). This suggests that StiX could act as a specificity factor for NixI, analogous to the specificity factors of type I restriction-modification systems (41), enhancing NixI-mediated cleavage at discrete regions.

Given StiX’s predicted homology to extracytoplasmic sigma factors, and because NixI cleaved DNA in the absence of StiX, we conjectured that StiX may enhance NixI-mediated cleavage by promoting NixI’s DNA-binding activity to specific sites. To assess whether StiX is a DNA binding protein, we performed electrophoretic mobility shift assays (EMSAs). To assay specificity, we used a probe that includes the *gp121* cut site identified *in vivo* during ICP1 infection (Figure 2C) and evaluated *in vitro* DNA binding with purified NixI+StiX. As a control, we included a probe from a region within *gp62* of ICP1, that has similar GC content to the *gp121* probe but was not identified as being cleaved *in vivo* and does not include the putative recognition motif. Despite predicted structural homology with DNA-binding proteins, StiX did not display any DNA-binding activity on its own, even at very high protein concentrations (Figure 4A and 4B). We observed NixI^N95A^ DNA binding at the highest protein concentration tested (100nM) for the *gp121* probe and minimal binding for the *gp62* probe (Figure 4, Supplemental Figure S7A-B), further supporting the specificity we observed in the nuclease assays (Figure 2E). StiX incubated with NixI^N95A^ showed a super-shift of the *gp121* probe, but not the control probe, suggesting that StiX may bind to NixI. There is a second similar motif (GAAAATCT) on the *gp121* probe that was also cut *in vivo* (Supplemental Figure S2) found ~60bp from the prominent cut site, opening the possibility that NixI may bind multiple sites on the *gp121* probe only when StiX is present. Of note, StiX appeared to enhance NixI binding activity at lower concentrations (50 and 25nM) (Figure 4A). Surprisingly, this was not apparent as a super-shift at these lower NixI^N95A^ concentrations, suggesting that perhaps StiX may interact transiently with NixI. Together, these data show that PLE-encoded NixI is a nuclease with intrinsic specificity for sites identified as being cleaved in ICP1, and the small downstream protein StiX enhances NixI-mediated cutting of target substrates possibly by increasing NixI’s DNA binding activity.

**Figure 4.**
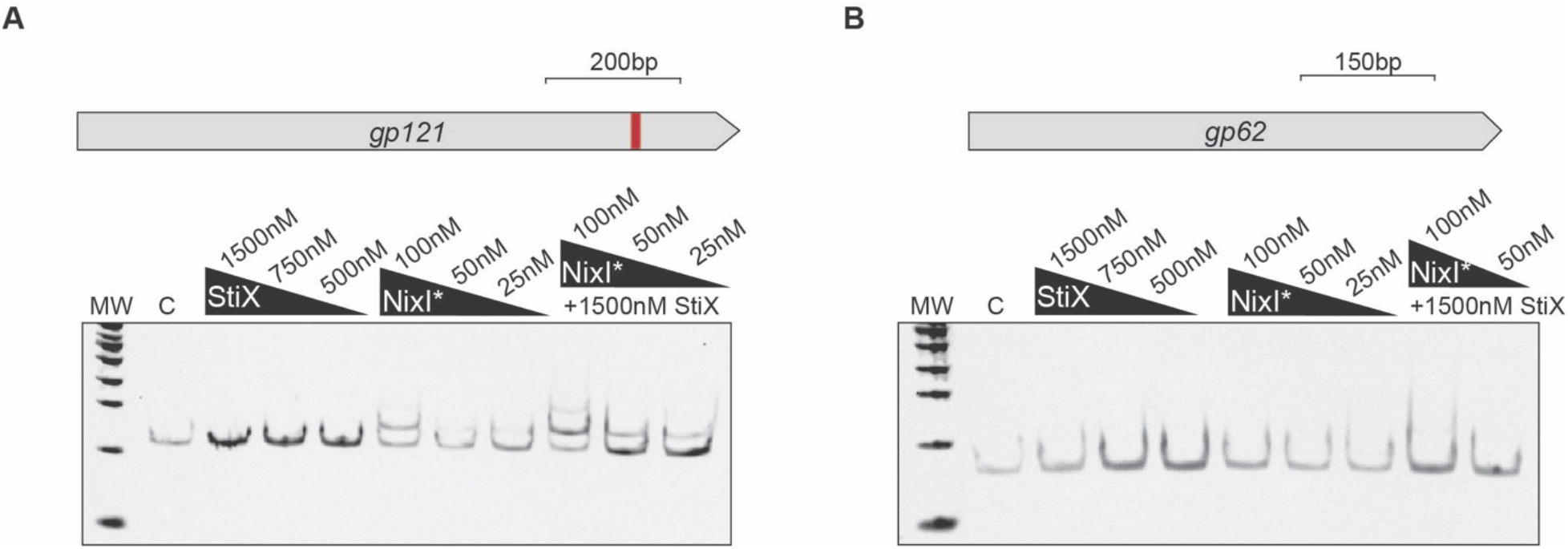
NixI and StiX bind specifically to a region of ICP1’s genome that is cut *in vivo* and *in vitro*. (A) Electrophoretic mobility shift assay (EMSA) using a probe spanning a 200bp region of *gp121* (top: the cut site identified *in vivo* shown as a red bar on the gene schematic), incubated with purified StiX or NixI^N95A^ either alone or in combination, as indicated by arrows and concentrations above the gels. (B) As in (A) but with a probe of similar length and GC content spanning a region in ICP1 not shown to be cut *in vivo*. C is no protein control and MW is molecular weight ladder. Replicate gels are shown in Supplemental Figure S7.

### NixI inhibits ICP1 progeny production and replication

Previous work has identified PLE-encoded mechanisms that effect ICP1 gene expression (13) and that collapse ICP1-induced lysis inhibition (15), however, none of these mechanisms alone are sufficient to prevent ICP1 progeny production, showing that there are additional mechanisms of PLE-mediated ICP1 interference. We anticipated that NixI cleavage of ICP1’s genome would have negative consequences for ICP1’s capacity to produce phage progeny. We first assayed whether *stiX* alone or *nixI*+*stiX* could inhibit plaque formation by ICP1 on a lawn of PLE(−) *V. cholerae* in the presence of inducer. Because of NixI’s toxicity to *V. cholerae* (Figure 2B) we were unable to assess plaque formation in a strain expressing NixI alone. Given that some toxicity is still observed in the presence of StiX (Figure 3C), we had to titrate down inducer until a visible lawn could form. Nonetheless, even at a low level of inducer (1/25^th^ of standard amount), we found that *nixI*+*stiX* inhibits ICP1 plaque formation ~100-fold (Supplementary Figure S8). Because NixI’s toxicity impaired our ability to perform plaque assays, we assessed *nixI*- and *nixI*+*stiX*-mediated inhibition of ICP1 using an efficiency of center of infection assay, which measures ICP1 virions produced from a single round of infection of *V. cholerae* by collecting virions produced from an infected host of interest and quantifying the progeny phage produced on a permissive host using a plaque assay (Figure 5A). *V. cholerae* expressing *nixI* or *nixI*+*stiX* reduced ICP1 progeny production 100-fold (Figure 5B). However, *nixI* was not required for PLE-mediated inhibition of ICP1, as PLE(+)Δ*nixI* was still inhibitory to ICP1 (Figure 5C). This corroborates previous work that showed no single PLE-encoded ORF, including *nixI*, is necessary for inhibition of ICP1 by plaque assay (15). NixI however is sufficient to limit ICP1 progeny production in a single round of infection, the first PLE-encoded product shown to recapitulate the block in progeny production from wild type PLE (Figure 5B and 5C).

**Figure 5.**
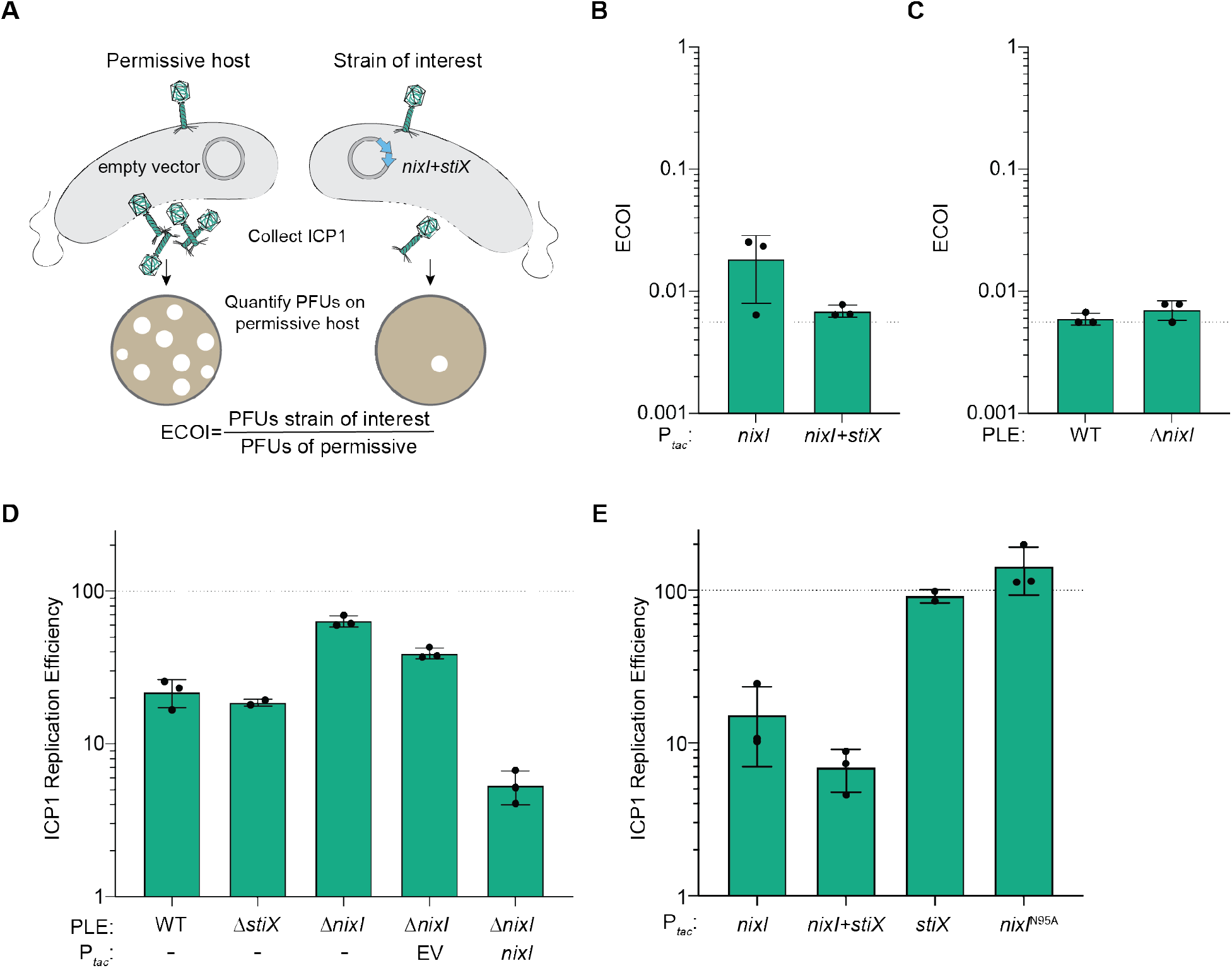
NixI and StiX are necessary and sufficient to inhibit ICP1 replication and sufficient to inhibit production of ICP1 progeny. (A) Schematic of efficiency of center of infection (ECOI) assay, where ICP1 produced by a single round of infection of a strain of interest (here, expressing *nixI*+*stiX*) is quantified as a ratio of the number of phage produced by a permissive host (left). (B) ECOI of ICP1 on PLE(−) *V. cholerae* expressing *nixI* or *nixI*+*stiX* from a plasmid, relative to phage progeny produced from an induced empty vector control. The dashed line indicates where the progeny phage output is equal to the input phage. (C) ECOI of ICP1 on PLE(+) *V. cholerae* or PLE(+)Δ*nixI* relative to phage progeny produced from a permissive control. The dashed line indicates where the progeny phage output is equal to the input phage. (D) ICP1 replication efficiency in the PLE(+) strains indicated relative to infection of permissive PLE(−) *V. cholerae* harboring an empty vector control (dashed line) 20 minutes post-infection as assessed by qPCR. (E) ICP1 replication efficiency of PLE(−) *V. cholerae* pre-induced to express the gene(s) indicated relative to infection of permissive PLE(−) *V. cholerae* harboring an empty vector control (dashed line) 20 minutes post-infection as assessed by qPCR. For B-E bar height displays mean, whiskers are standard deviation, and dots indicate independent biological replicates.

We anticipated that *nixI*+*stiX* inhibits ICP1 progeny production by impairing phage genome replication, which is a hallmark of PLE activity (18). We therefore measured ICP1 genome replication by qPCR in a wild type PLE(+) host compared to Δ*stiX* and Δ*nixI* mutant derivatives. As was previously observed (18), PLE reduced ICP1 genome copy to ~20% of the levels observed in a permissive infection, and although StiX enhanced NixI activity *in vitro* (Figure 3G), we found that *stiX* is not required for PLE-mediated inhibition of ICP1 DNA replication (Figure 5D). However, deletion of *nixI* restored ICP1 replication to ~65% of the levels of replication observed in a permissive host (Figure 5D). Complementation of *nixI in trans* reduced ICP1 replication to a greater extent than wild type PLE (Figure 5D). This is likely because endogenous *nixI* is not expressed until mid-way through infection, between 8 and 12 minutes post-infection (37). At this point in infection, ICP1’s has undergone several rounds of theta replication and is transitioning to rolling circle replication, while PLE replication begins to accelerate (18). Pre-induction of *nixI* as was done here (Figure 5D) would likely interfere with all stages of ICP1 replication and could therefore exceed PLE’s impact on ICP1 DNA replication. To assess if NixI or Nix+StiX directly impact ICP1’s genome replication in the absence of other PLE-encoded products, we also evaluated ICP1 genome replication in a PLE(−) background. As seen with *nixI* complementation *in trans*, ectopic expression of *nixI* and *nixI*+*stiX* was sufficient to inhibit ICP1 replication to 10% that of a permissive infection, again greater inhibition than is observed in the context of wild type PLE (Figure 5E). Induction of *stiX* or the catalytically dead *nixI*^N95A^ did not inhibit ICP1 replication (Figure 5E), supporting that catalytic activity of NixI is necessary to inhibit ICP1 replication. Together these data demonstrate that *nixI* is a potent inhibitor of ICP1 replication, resulting in decreased virion production.

### NixI and StiX promote PLE replication

PLE replication relies on hijacking replication resources from ICP1, leading us to hypothesize that PLE may limit ICP1 replication to successfully co-opt replication machinery for its own replication. This suggested that NixI-mediated cleavage of ICP1’s genome and its subsequent limitation of ICP1 replication may be necessary for PLE to replicate efficiently. We therefore assayed PLE replication efficiency of PLE Δ*nixI* relative to wild type PLE by qPCR (Figure 6A). During ICP1 infection we observed that PLE replication is reduced to 25% of wild type in absence of *nixI* (Figure 6A). Complementation of *nixI in trans* restored PLE replication to wild type levels. Importantly, these data help us to contextualize the effect of *nixI* expression on ICP1 replication. Because *nixI* induction is toxic to *V. cholerae* (Figure 2B), we initially had concerns that the observed reduction in ICP1 replication in the presence of *nixI* was the result of indirect effects on *V. cholerae*, rendering the host unable to support phage infection and DNA replication in general. However, PLE replication is increased by *nixI* expression *in trans* (Figure 6A), demonstrating that *nixI*-mediated inhibition of ICP1 replication (Figure 5E) is specific and not a pleiotropic effect of cellular toxicity. Further, we observed that PLE replication is similarly reduced to 25% of wild type in the absence of *stiX* (Figure 6A). In contrast, ICP1 replication was not restored during infection of PLE(+)Δs*tiX*, supporting the model that StiX functions to restrict non-specific cleavage by NixI, perhaps helping to protect PLE from cleavage.

**Figure 6.**
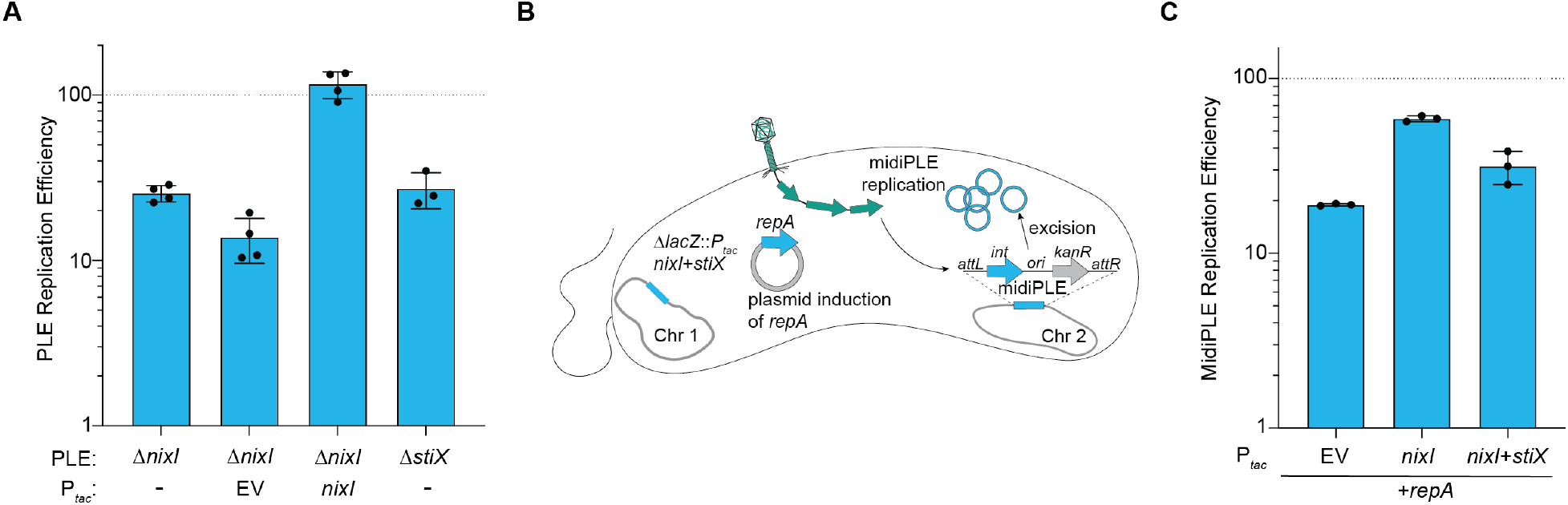
Expression of *nixI* is required for PLE replication and can augment replication of a minimal replicating PLE (midiPLE) (A) Replication efficiency of *V. cholerae* PLE mutant and complemented derivatives 20 minutes post-infection by ICP1 relative to replication of wild type PLE (dashed line). (B) Model of midiPLE induction and replication during ICP1 infection. Plasmid-encoded *repA*, PLE’s essential replication protein(18) and chromosomally encoded *nixI*, *nixI*+*stiX* or an empty vector were pre-induced for 15 minutes prior to infection by ICP1. ICP1 infection triggers midiPLE excision from the *V. cholerae* chromosome and replication of midiPLE is detected by qPCR. (C) Replication efficiency of midiPLE relative to wild type PLE (dashed line) 20 minutes post-infection. For A and C, bar height indicates mean, whiskers are standard deviation, dots are independent biological replicates.

Previous work characterizing PLE replication identified PLE’s minimal unit of replication, (referred to as midiPLE) comprised of PLE’s attachment sites, integrase, and origin of replication integrated into PLE’s native locus in *V. cholerae* (Figure 6B). When ICP1 infects, midiPLE excises from the chromosome, and in the presence of PLE’s essential replication initiation factor, *repA*, midiPLE replicates to high levels; however, midiPLE replication does not reach the same robust levels of replication as is observed with the full length wild type PLE (18). As demonstrated above, NixI is required for wild type levels of PLE replication (Figure 6A), therefore we asked whether NixI +/− StiX could boost midiPLE replication to levels more comparable to wild type PLE. Induction of *nixI* increased midiPLE replication to ~60% of wild type PLE replication, greater than *repA* driven midiPLE replication alone (Figure 6C). Interestingly, *nixI*+*stiX* together did not boost midiPLE replication beyond that of *nixI* alone (Figure 6C). The boost in midiPLE replication due to NixI expression was surprising, as we anticipated NixI expression could have hindered midiPLE replication due to cleavage of *V. cholerae*’s chromosome before midiPLE has excised. The impact of NixI on midiPLE replication could be due to inhibition of ICP1, or NixI may have a direct role in PLE replication, as nicking enzymes are required factors for initiating RCR of plasmids and viral genomes (23). Since we anticipate that PLE also undergoes a transition to RCR to linearize its genome for packaging, NixI may act as the nicking nuclease for initiating PLE RCR, which is often the same region where packaging is initiated from (the *pac* site). To test whether NixI can cleave PLE’s RCR ori, we identified a candidate region in PLEs ORF-less space to clone onto our plasmid as a substrate for nuclease assays. We reasoned that PLE’s RCR ori would be highly conserved among all known PLEs, and cloned a 500bp region previously identified as highly conserved (21) (Supplemental Figure S9A). We found that even at high protein concentrations NixI exhibited minimal cleavage of this substrate suggesting NixI does not directly cut PLE’s RCR ori (Supplementary Figure S9B). Additionally, if NixI directly cuts midiPLE’s RCR origin, pre-induction of *nixI* would be detrimental to midiPLE replication, as it would nick the origin potentially before the expression of the remaining proteins required for RCR (such as single stranded-binding protein) and midiPLE would likely fail to replicate. Because NixI increases midiPLE replication (Figure 6C), even when ectopically expressed prior to infection, we anticipate NixI is not PLEs RCR initiator nickase, but rather cleaves ICP1’s genome to liberate replication resources for midiPLE. The targeted nicking of ICP1 could liberate replication resources for PLE and hinder ICP1’s ability to package full-length genomes. Although the difference is not significant, *stiX* expression in concert with *nixI* further reduced ICP1 replication and progeny formation (Figure 5B and 5E) but did not confer the boost in midiPLE replication seen with *nixI* alone (Figure 6C).

### Homologs of NixI are a conserved feature of an expanded family of phage satellites in Vibrios

To date, PLEs have only been found in toxigenic *V. cholerae* and have been shown to be dependent on ICP1. Accordingly, all PLE(+) *V. cholerae* that have been isolated possess the lipopolysaccharide O1 antigen that serves as the receptor for ICP1(43). Bioinformatic searches for PLEs in sequenced genomes have searched for nucleotide similarity, as PLEs 1-5 all share homology at the nucleotide level. In an effort to predict NixI’s domain organization and function, we used HHblits (44) to look for homologs of NixI^PLE 1^ and found proteins that share ~25% amino acid identity in non-toxigenic *V. cholerae* (e-value=0.0000072). Notably, NixI homologs outside of those found in PLEs in toxigenic *V. cholerae* do not share detectable homology at the nucleotide level. Further investigation of the genomic context of these NixI homologs identified an expanded family of putative phage parasites similar to PLEs, as we elaborate on below. To determine whether the satellites from non-toxigenic *V. cholerae* were indicative of an even larger family of parasites, PSI-BLAST searches were performed with the NixI homologs from non-toxigenic *V. cholerae* PLEs and excluding *V. cholerae* as a subject. This search returned several homologs in other Vibrios such as *V. parahaemolyticus* and *V. casei*. To expand this family of potential PLEs, reciprocal searches were performed with the tyrosine recombinase/integrase from *V. parahaemolyticus* strain S042 and additional elements were gathered. In all cases, the NixI homologs were found on putative mobile genetic elements which possess key features of PLEs. First, they all encode a 5’ integrase/recombinase; either a large serine recombinase (which share ~50% amino acid identity with PLE-encoded large serine recombinases from toxigenic *V. cholerae*) or a tyrosine recombinase (Figure 7A). All putative parasites are ~17kB in length and, like PLEs from toxigenic *V. cholerae*, encode for almost exclusively hypothetical proteins. However, PFAM domain searches indicated that the satellites have ORFs with domains in previously characterized PLE ORFs (Figure 7A). Specifically, we searched for proteins with homology or shared domains with PLE 1’s CapR (T5ORF172 domain) (13), RepA (RepA_N domain) (18), StiX (sigma factor like), and LidI (transmembrane domain) (15). All putative satellites encode a predicted replication initiation protein at the same location as PLEs′ RepA, but interestingly, we saw two different replication proteins: either an N-terminal RepA_N domain or a C-terminal DNA Pol I domain. The putative parasites have a small transmembrane domain protein similar to PLE’s LidI (15), located towards the 3’ end, suggesting that disrupting lysis inhibition may be a conserved function of phage parasites found in other *Vibrio* species. None of the parasites possessed a protein with a T5ORF172-domain protein. However, we noted the occasional occurrence of Bro-N domain containing proteins, a DNA-binding domain often associated with T5ORF172 domains (45). In addition, all of the putative parasites are syntenic with PLEs, possessing a large ORF-less space where PLEs’ origin of replication (18) and putative *pac* site are located, as well as a smaller ORF-less space upstream of *nixI*, where PLEs encode a predicted small RNA (37). Of note, none of the putative satellites possessed an identifiable homolog of StiX, as the gene immediately 3’ of *nixI* shared a predicted peptidase function as indicated by PFAM domain searches (Figure 7A). This suggests that although *nixI* may play a conserved function for an expanded family of phage parasites, PLEs in toxigenic *V. cholerae* appear to be unique in possessing *stiX*, which we have shown here modulates *nixI* activity.

**Figure 7.**
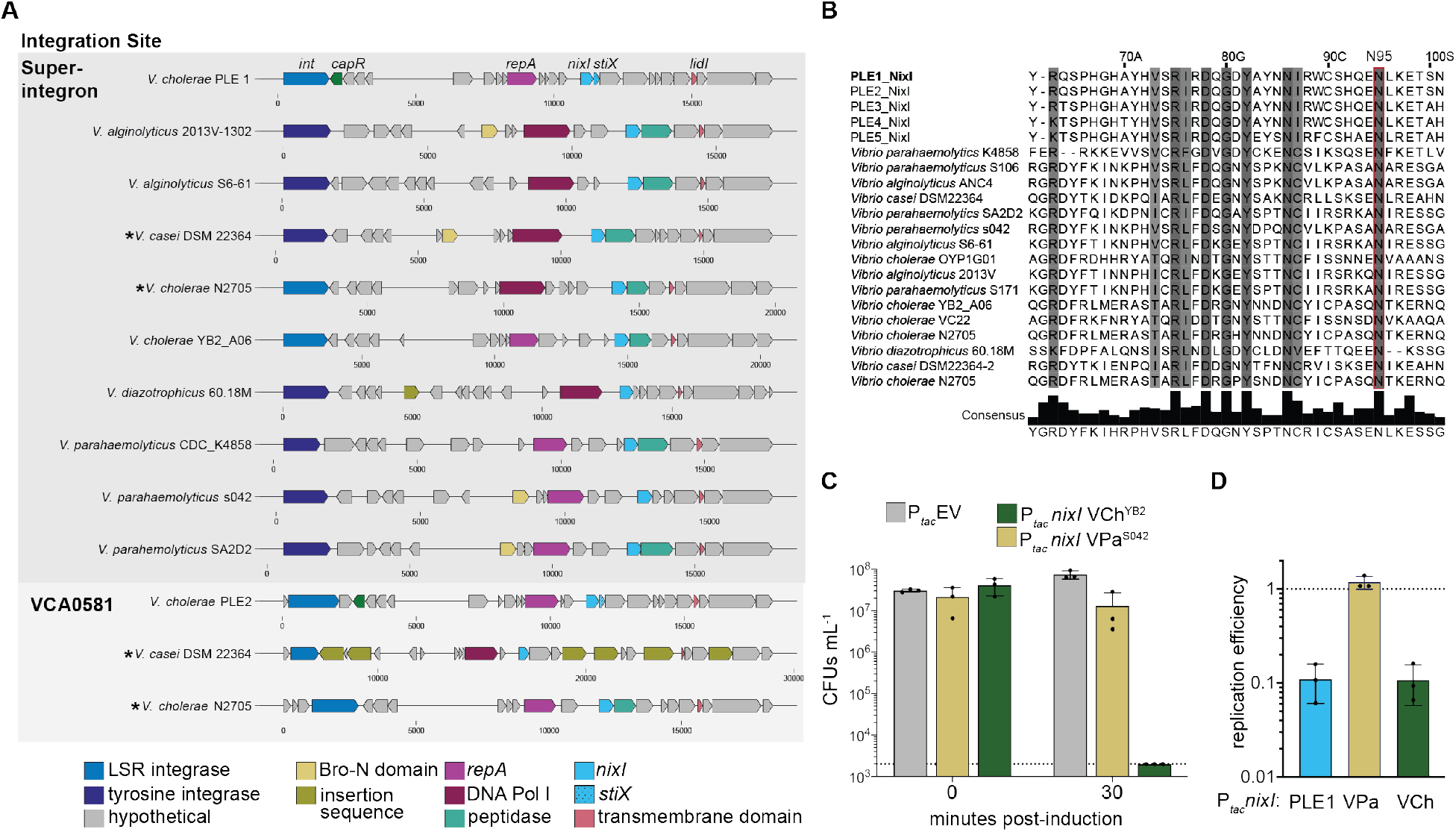
NixI homologs characterize an expanded family of putative phage parasites in *Vibrio* species. (A) Gene maps (scale below cartoon genes displays relative nucleotide position) of putative phage parasites from Vibrios, where homologs of functionally characterized genes from PLE 1 (labeled top) are colored according to shared predicted function identified by PFAM domains or HHPred. Top: phage parasites integrated into the repeat regions of the superintegron, bottom: phage satellites integrated into the PLE 2 locus (*vca0581*). Asterisks are used to note where two similar phage satellites occur in different locations in the same genome. LSR = large serine recombinase. (B) MUSCLE alignment of NixI homologs, showing PLE 1’s experimentally determined catalytic asparagine boxed in red (N95), and residues sharing at least 70% consensus shaded in grey. (C) Cell viability as measured by CFUs ml^−1^ of *V. cholerae* expressing *nixI* homologs from *V. cholerae* YB2_A06 (VCh^YB2^) and *V. parahaemolyticus* s042 (VPa^s042^), plated prior to induction and 30 minutes post-induction. The dashed line indicates the limit of detection. (D) Same colors as in C, showing ICP1 replication efficiency relative to *V. cholerae* harboring an empty vector control (dashed line) 20 minutes post-infection as assessed by qPCR, pre-induced to express *nixI* homologs. For C and D, bar height displays mean, whiskers are standard deviation and dots indicate independent biological replicates

We next looked at the sequence flanking the putative satellites to determine if they integrate into conserved attachment sites in their bacterial hosts. In the laboratory, PLEs 1, 3, 4 and 5 from toxigenic *V. cholerae* integrate into the superintegron, in *V. cholerae* repeats (VCRs) that occur >100 times, each serving as a potential attachment site (17). Although long-read sequencing is required to resolve the repetitive nature of the superintegron to definitively state where these PLEs integrate, the majority of satellites found in this expanded family are integrated into the superintegron repeats (46), even though some possess a tyrosine recombinase unrelated to the large serine recombinases found in PLEs from toxigenic *V. cholerae*. PLE 2 is unique in integrating into the M48 metallopeptidase gene (*vca0581*) outside of the superintegron (17). Strikingly, two of the identified *Vibrio* genomes, *V. cholerae* N2705 and *V. casei* DSM22364, have multiple PLEs: one integrated into a superintegron repeat and a second in the PLE 2 locus. Multiple PLEs in a single *V. cholerae* genome has not previously been described. The presence of multiple satellites similar to PLEs found in the same genome suggests the modular nature of PLEs that has been observed (17) could be driven by recombination between PLEs in the same host. The second putative satellite in *V. casei* (in the PLE 2 locus) is littered with insertion sequences, suggesting that there may be no fitness benefit to having multiple phage satellites and that they can be lost through degeneration (Figure 7A).

NixI homologs from PLEs in toxigenic *V. cholerae* are all highly similar, sharing >65% amino acid identity. In contrast, NixI homologs from the putative satellites outside of toxigenic *V. cholerae* are highly divergent, sharing only 20-25% amino acid identity. A MUSCLE alignment of all unique NixI homologs from *Vibrio* species indicates that the HNH/N catalytic asparagine identified in this study is conserved (Figure 7B). To determine if NixI is functionally conserved among the newly identified phage satellites, we expressed *nixI* from *V. cholerae* YB2_A06 (an environmental non-toxigenic isolate) and from *V. parahemolyticus* S042 in our lab strain of toxigenic *V. cholerae*, and the impact of NixI induction on cell viability was assayed. Induction of the homolog from *V. cholerae* YB2_A06 completely inhibited colony formation of our toxigenic laboratory strain of *V. cholerae*, while the homolog from *V. parahaemolyticus* was only mildly toxic (Figure 7C). To determine if these *nixI* homologs are inhibitory to ICP1, we quantified ICP1 replication in *V. cholerae* pre-induced to express either homolog, or *nixI* from PLE 1, and compared replication efficiency to that of a permissive host with the empty vector. Interestingly, the homolog from *V. cholerae* YB2_A06 inhibited ICP1 replication, yet the homolog from *V. parahaemolyticus* did not (Figure 7D). *V. cholerae* YB2_A06 is a non-toxigenic non-O1 isolate that would not be susceptible to ICP1 infection. Induction of PLE’s genetic program requires infection by ICP1 (37), so specificity of NixI homologs for their cognate phages could be driven by expression only in response to infection by their viral host, rather than recognition of a motif unique to their host phage’s genome. Together, we show that PLEs are not restricted to toxigenic *V. cholerae*, but functionally analogous elements also reside in non-toxigenic *V. cholerae* as well as other Vibrios. Although members of this expanded family of phage satellites remain to be experimentally validated, these data suggest that PLE-encoded strategies that restrict ICP1, such as *nixI*-mediated cleavage, may be conserved among divergent satellites and represent a mechanism to parasitize a broader class of inducing phage.

## Discussion

Phage parasites can be considered host defense systems, as the parasite inhibits production of its viral host, benefiting the bacterial population (4, 17). However, phage parasites are fundamentally selfish elements, promoting both their *vertical* transmission by protecting the bacterial host population from viral takeover, as well as their own *horizontal* transmission through parasitism of their viral host’s structural components (7, 13, 47). Phage parasites employ finely tuned mechanisms to prevent packaging of the viral host’s genome, but allow for progression of the viral host’s gene expression program for production of structural proteins which are hijacked by the phage parasite. For this reason, it is perhaps unsurprising that all phage parasites studied outside of *V. cholerae* reduce but do not eliminate production of their viral host (7, 9, 48, 49). *V. cholerae*’s PLEs are unique in their capacity to completely restrict their viral host ICP1 (17), yet, the PLE-encoded mechanisms responsible for ICP1 restriction have remained mysterious. In this work, we show that NixI from PLE 1 functions as a nuclease effector, adopting a strategy that is widely used by bacterial defense systems to restrict phage production (50–52). We propose a model where NixI functions as a nicking endonuclease, introducing site-specific nicks into ICP1’s genome. Anti-phage defense systems that encode nicking endonucleases have recently been characterized suggesting nicking is an effective strategy to destroy invading viral DNA (32, 53). We show here that NixI is the most inhibitory PLE-encoded product characterized thus far, inhibiting ICP1 progeny production to the same extent as is seen with PLE, and reducing ICP1 replication by an order of magnitude. Targeting viral host replication with a nuclease effector is a strategy of viral host interference that, to our knowledge, has not been previously reported for phage parasites. Though as we discuss below, PLE-mediated restriction of ICP1 replication through NixI does not completely prevent ICP1 replication, as PLE relies on the maintenance of ICP1’s genome for expression of replication (22) and structural (13) proteins that PLE parasitizes. NixI is not necessary for PLE-mediated inhibition of ICP1, consistent with there being multiple PLE-encoded mechanisms to restrict ICP1 (15). While the remaining mechanisms of ICP1 restriction remain to be determined, they may target nodes in ICP1s lifecycle to divert structural components towards PLE packaging; a strategy used by all known phage parasites studied to date (11, 54, 55).

Several lines of evidence implicate NixI as the PLE-encoded factor that inhibits ICP1’s transition to RCR. First, bioinformatic analyses identified NixI as the only putative nuclease encoded by PLE. Additionally, transcription of the *nixI* operon peaks between 12 and 16 minutes post-infection, at a time when ICP1 is transitioning to RCR (18, 37). Additionally, NixI is necessary for the appearance of cut sites in ICP1’s genome, which are evident at 16 minutes post-infection of PLE(+) *V. cholerae*. PLE-mediated inhibition of ICP1’s RCR would prevent ICP1 from linearizing its genome, which is necessary for efficient packaging. In this way, inhibition of RCR may be functionally analogous to a strategy used by phage parasites in *Staphylococcus aureus* (known as SaPIs) to prevent packaging of their viral host’s genome. SaPIs inhibit their viral hosts’ small terminase (TerS) subunit and encode their own replacement (56). TerS is the component of the packaging machinery that specifically recognizes the *pac* site on the viral DNA to initiate translocation of the viral genome into virions (26). By blocking TerS and encoding their own, SaPIs ensure that their own genome is preferentially packaged. Adopting a similar strategy, divergent satellites found in *E. coli* encode a protein that redirects the viral host’s TerS to specifically package the parasites’ genome (57). Our analyses suggest that ICP1 packaging is still inhibited by PLEΔ*nixI*, as cleavage of ICP1’s *pac* site is still blocked (Supplemental Figure S10). This suggests that PLE encodes additional inhibitory mechanism(s) that may block ICP1’s terminase complex, and may act synergistically with NixI to ensure ICP1’s genome is not packaged.

We show here that NixI nuclease activity is augmented by StiX, which reduces *nixI* toxicity when co-expressed in *V. cholerae*. We propose that StiX restricts NixI nuclease activity by aiding its cleavage of specific motifs, which could allow survival of *V. cholerae* if it is able to repair a limited number of nicks compared to widespread cleavage by an undirected NixI. However, it is possible that ectopic expression of *nixI* and *stiX* do not recapitulate native expression or stoichiometry, and further work should address whether native protein levels are comparable to those used *in vitro* in this study. Although we attempted to address this by quantifying tagged NixI produced from its native context, we were unable to clone an affinity tag into NixI in its native PLE context, and our attempts to procure an antibody for NixI from a commercial source were unsuccessful due to NixI’s toxicity. Regardless of how StiX alters NixI binding, we propose that it functions to protect PLE from self-cleavage. An essential component of nuclease effectors is the ability to discriminate self from non-self. In support of this, PLE replication is not inhibited, but is actually enhanced, by ectopic expression of *nixI* and *stiX*. Additionally, PLE replication is reduced in the absence of StiX, which would be expected if StiX functioned to protect PLE from NixI cleavage. Initially we hypothesized PLE 1 was protected from NixI activity because it lacks the consensus motif identified in ICP1 (Figure 1D). However, PLE 2 encodes for identical NixI and StiX proteins, yet contains three occurrences of the predicted recognition motif. PLE 2 replicates during ICP1 infection (17) and shows the same temporal *nixI*+*stiX* expression pattern as PLE 1 (37) suggesting that PLE 2 is not cleaved by NixI. Together these data suggest that the motif alone is not sufficient to determine cutting. We propose that DNA topology may be an additional component of NixI+StiX specificity, as only supercoiled plasmids were cleaved by NixI (Figure 2 and 3, Supplemental Figure S5), yet our EMSAs show that NixI+StiX can still bind linear DNA *in vitro*. Further work is needed to determine how PLE is protected from NixI-mediated cleavage.

Established and novel phage defense systems can be predicted *in silico* through the presence of conserved domains or by their genomic proximity to known defense systems (44). Our work complements recent work showing that phage parasites are likely ubiquitous in bacterial genomes (11, 12) and highly similar parasites can be found in divergent taxa, such as the parasite P4 that was previously thought to be restricted to *Escherichia coli* but was recently found across several genera of Enterobacterales. (58). Phage parasites lack hallmark genes allowing for easy identification but can be identified as mobile genetic elements carrying an integrase/recombinase. Additional gene cargo carried by phage parasites can influence the fitness of their bacterial hosts, beyond their established role in restricting viral host propagation, as new phage parasites have been shown to encode defense systems that function more broadly against infecting phages (59). Demonstrating bona fide phage parasitism is challenging given that a compatible tripartite system must be identified. While the cellular host is known from the integrated parasite genome, the cognate viral host may not be as obvious, especially if the viral host is not temperate. Interestingly, phages similar to ICP1 (60) have been isolated and shown to infect non-cholera Vibrios (61) which may be candidate viral hosts for the new parasites identified here. To identify the viral host, searching for homologs of phage parasite encoded proteins can help find candidate hosts. Previous work has shown that PLEs from toxigenic *V. cholerae* have domesticated a homing endonuclease from ICP1 to manipulate ICP1 gene expression, showing that parasites can coopt genes from their viral hosts (13). Interestingly, we found homology between the Bro-N domain containing proteins from *V. parahaemolyticus* and *alginolyticus* and a Bro-N domain protein from the Vibrio phage 11895-B1 (HQ634157.1), suggesting this phage as a candidate viral host for these putative satellites. Complicating the identification of viral hosts is the arms race that can occur between viral hosts and their parasites, as occurs with ICP1 and PLEs, which may obscure inhibitory activity if the phage being tested has evolved to antagonize the phage parasite in question (14, 21). The outcome of phage infection can be hard to predict based solely on the genome sequences of bacterial hosts and their phages, influencing microbial ecology and complicating phage therapy efforts. Vibrios are responsible for diseases in aquaculture, prompting research into the use of phage therapy to control outbreaks (62). Phage parasites may hinder these efforts, and research into the viral host range and prevalence of putative parasites in aquaculture settings should be considered before application of phages.

Together we have shown that PLE encodes a nuclease and specificity protein that specifically targets ICP1’s genome for cleavage, culminating in decreased ICP1 replication and inhibition of ICP1 progeny production. Although NixI homologs are a feature of an expanded family of putative phage satellites, StiX is unique to PLEs in toxigenic *V. cholerae*. NixI and StiX are expressed specifically in response to ICP1 infection, aiding in PLE’s efficient cooption of ICP1 components for its own lifecycle. This work furthers our understanding of the tight co-evolutionary relationship between ICP1 and PLE, where adaptations and counter-adaptations between these two genomes have been ongoing for nearly 100 years, shaping phage infection dynamics within cholera-infected patients.

## Data Availability

Prior to publication, the sequencing data from phage infected *V. cholerae* PLE(+)Δ*nixI* generated in this study will be deposited in the Sequence Read Archive database under accession codes TBD.

## Acknowledgements

We would like to thank all current and former members of the Seed Lab for their many helpful conversations and feedback. Specifically, we would like to thank Dr. Angus Angermeyer for his help analyzing ICP1’s coverage ratios. We would also like to thank the Komeili lab for use of their AKTA for protein purification and the University of California, Berkeley QB3 Core Facility for assistance with whole genome sequencing.

## Funding

This work was supported by the National Institute of Allergy and Infectious Diseases (grants R01AI127652 and R01AI153303 to K.D.S.). K.D.S. is a Chan Zuckerberg Biohub Investigator and holds an Investigators in the Pathogenesis of Infectious Disease Award from the Burroughs Wellcome Fund. K.N.L. was supported by an NSF Graduate Research Fellowship (Fellow ID no. 2017242013). P.D. was supported by a Rose Hills Summer Undergraduate Research Fellowship.

## Conflict of Interest Disclosure Statement

K.D.S. is a scientific advisor for Nextbiotics, Inc.

