## Supplemental tables and figures for "A phage parasite deploys a nicking nuclease effector to inhibit replication of its viral host"

### Supplementary Figures

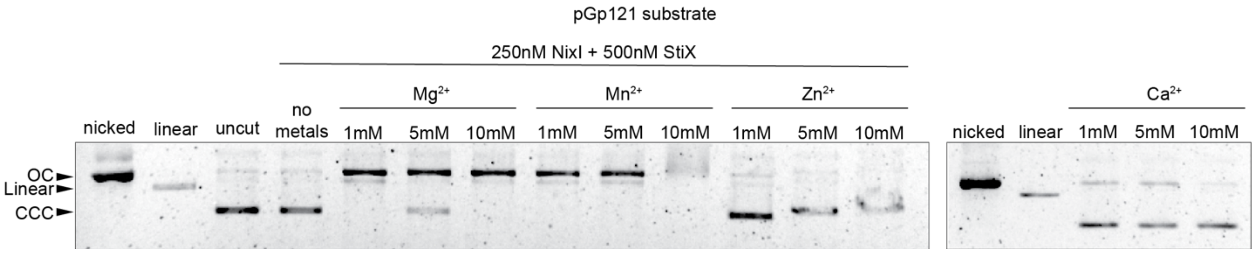


**Figure S1. NixI divalent metal requirements.** (A) NixI (250nM) and StiX (500nM) were incubated for 15 minutes with pGp121 substrate and cutting was tested with decreasing concentrations of $Mg^{2+}$, $Mn^{2+}$, $Zn^{2+}$ and $Ca^{2+}$ compared to buffer without metals, and compared to a nicked and linear plasmid control. Arrows on the left indicate covalently closed circular DNA (CCC), linear DNA or open circular (OC) DNA (nicked DNA).

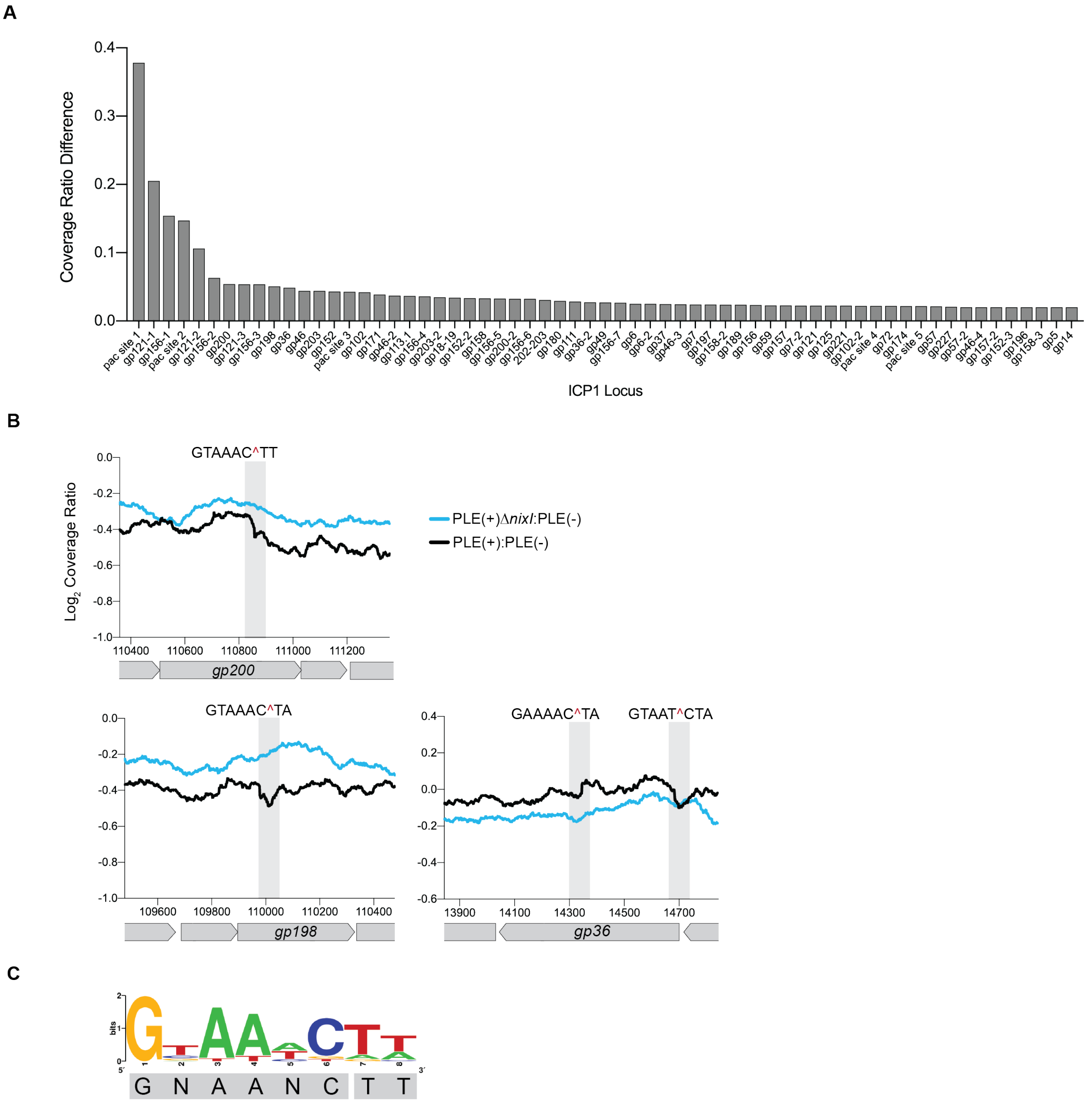


**Figure S2. NixI cleaves additional putative cut sites from ICP1** (A) $Log_2$ coverage ratios PLE(+)/PLE(-) of additional coverage drops in ICP1's genome with a difference in nucleotide coverage >$log_2$(0.026). (B) $Log_2$ coverage ratios PLE(+)/PLE(-) and PLE(+)Δ*nixI*/PLE(-) of the next three most dominant coverage drops in ICP1's genome, showing that the coverage drops are abolished in a PLE(+)Δ*nixI* host. (C) The consensus motif produced from 16 unique sites in ICP1's genome that are cleaved only during infection of a PLE(+) host. In regions with multiple cleavage sites within 100bp of each other, only the most dominant site was used to generate the motif.

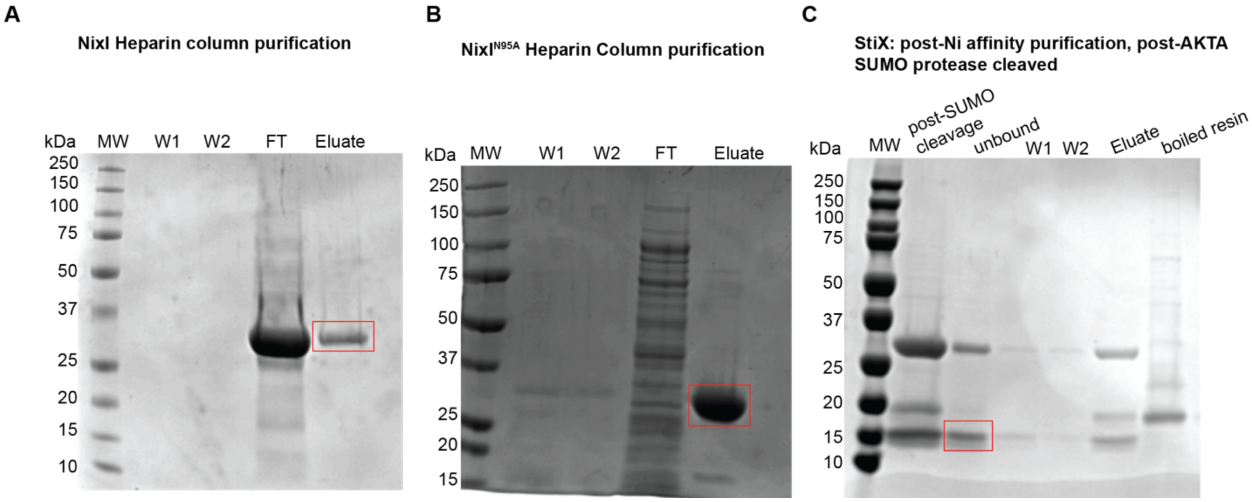


**Figure S3. Protein gels of purified proteins used in this study.** (A) Wild type NixI after purification with a heparin column, where eluate was used for the assays in this study (B) NixI$^{N95A}$ purification with a heparin column, where the eluate was used for the assays in this study. (C) StiX after cleavage of the SUMO tag (input shown in lane 2) and batch purification with Ni-NTA resin to remove free SUMO protease and cleaved tag, where the unbound protein was used in the assays in this study. MW=molecular weight ladder (in Kilodaltons, W1=wash 1, W2=wash 2, FT=flow through/unbound protein. Red boxes highlight the purified protein of interest.

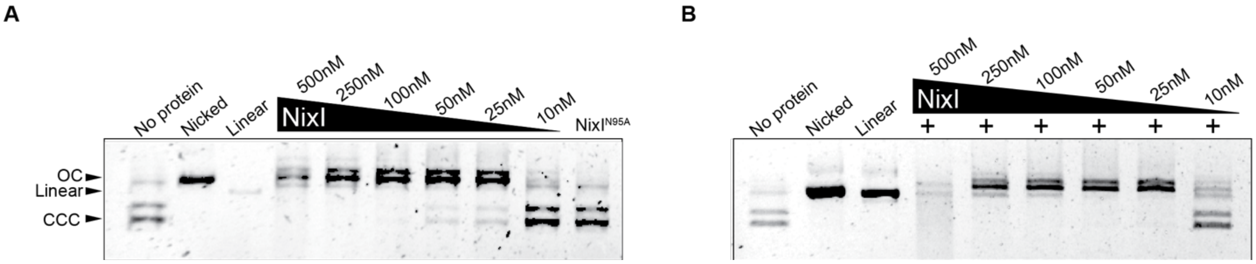


**Figure S4. NixI cleaves additional ICP1 cut sites *in vitro*.** Left: cutting by purified NixI or NixI$^{N95A}$ of a plasmid containing 500 basepairs flanking ICP1's cut region in *gp156*, arrows indicating supercoiled plasmid (CCC), linear plasmid and nicked (OC) plasmid. Right: NixI supplemented with 500nM of StiX shows enhanced cleavage of the plasmid with *gp156*.

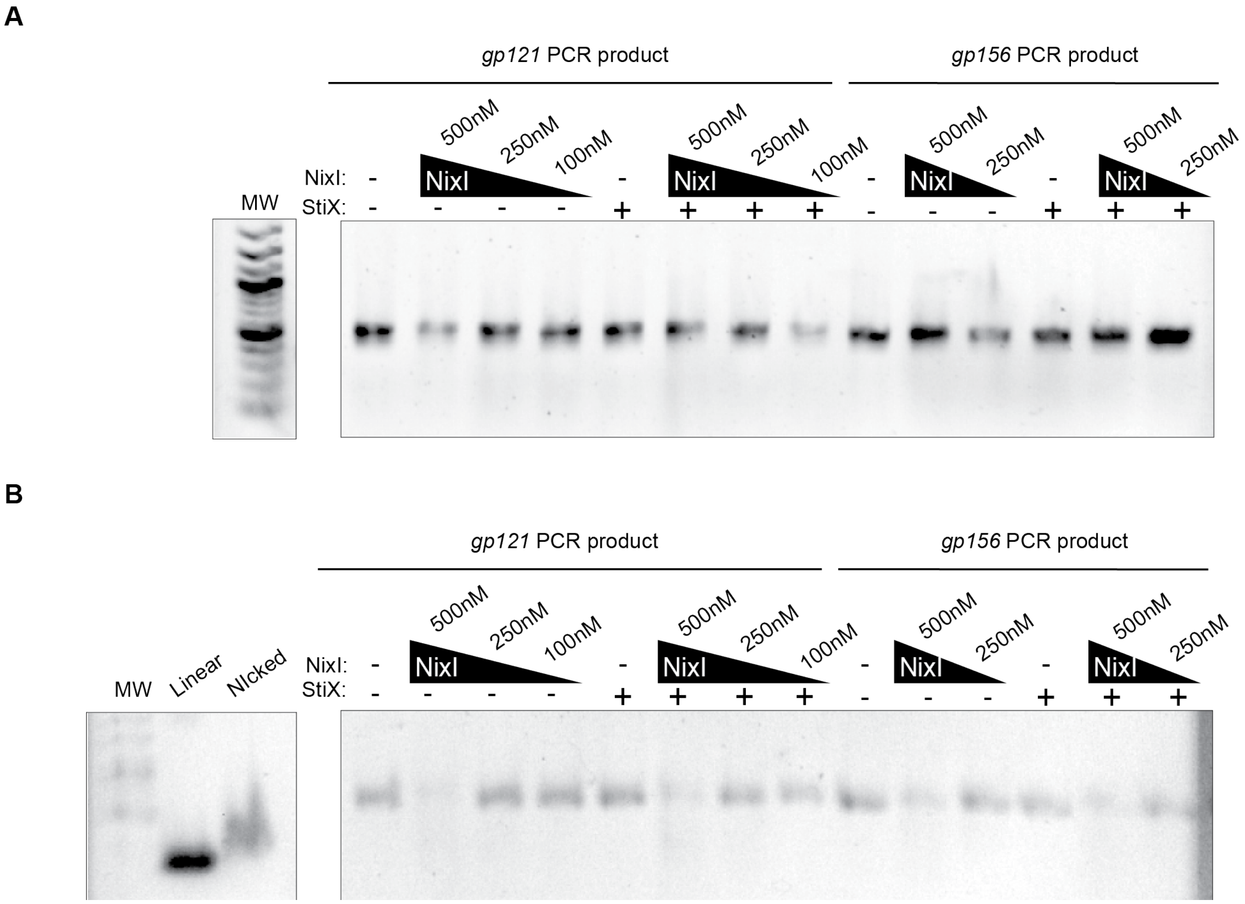


**Figure S5. NixI cannot cleave PCR products spanning cut regions from ICP1.** (A) Native gels of PCR products incubated with decreasing concentrations of NixI and a constant concentration of StiX (500nM) with substrates spanning *gp121* and *gp156* cut regions which were used as the insert in the plasmids as in Figure 2C and Supplementary Figure S4. (B) Alkaline denaturing agarose gel of the same reactions from A, with a nicked PCR product control and linear uncut control on the far left.

A

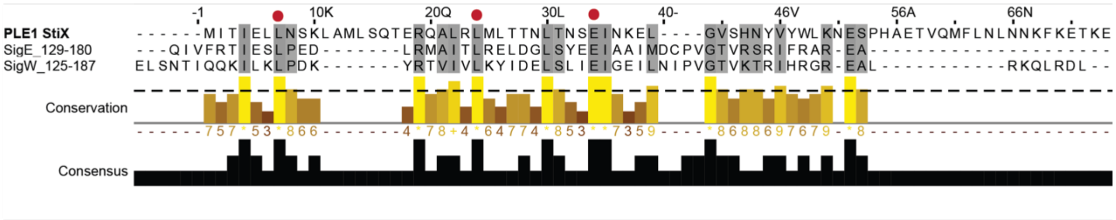


B

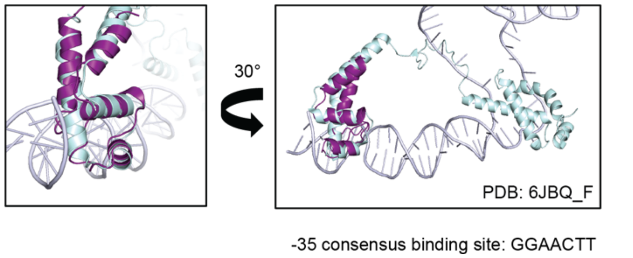


**Figure S6. StiX shares predicted structural homology with alternative sigma factors SigE (*Escherichia coli*) and SigW (*Bacillus subtilis*).** (A) MUSCLE alignment of Stix, SigE and SigW showing conserved residues required for sigma factor structural conformation in red circles, where grey shading highlights residues that are >65% conserved. Residues that contact DNA basepairs and determine sequence specificity are variable and not conserved. (B) A predicted structure of StiX generated by i-TASSER overlaid on the $\sigma^4$ domain of SigE, which contacts the -35 site and recognizes the consensus sequence GGAACTT, rotated 30° to show contacts with DNA.

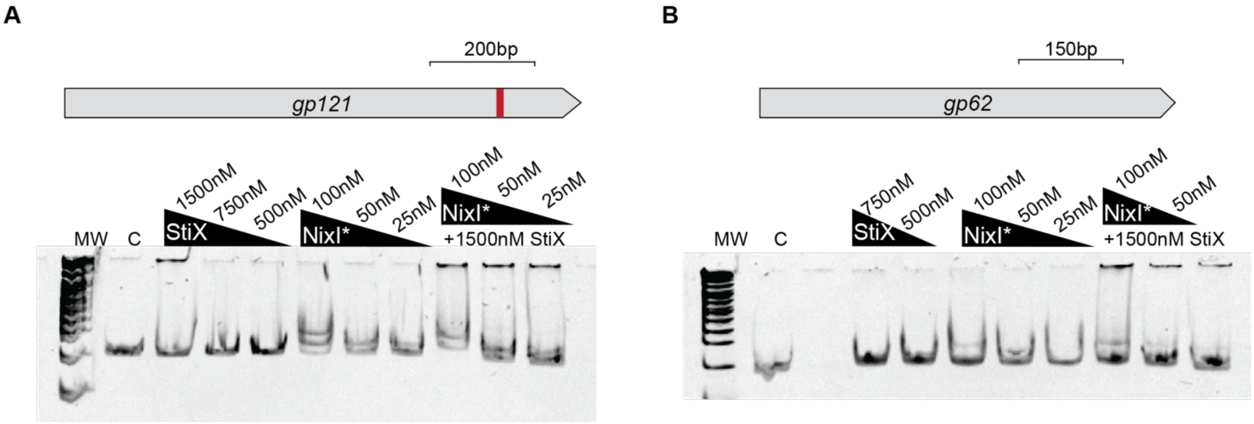


**Figure S7. Replicate electrophoretic mobility shift assays (EMSAs)** StiX, NixI$^{N95A}$ or both proteins together incubated with a probe spanning *gp121* (in A) or *gp62* (in B), performed as in Figure 4, shows enhanced binding of *gp121* with NixI$^{N95A}$ with StiX.

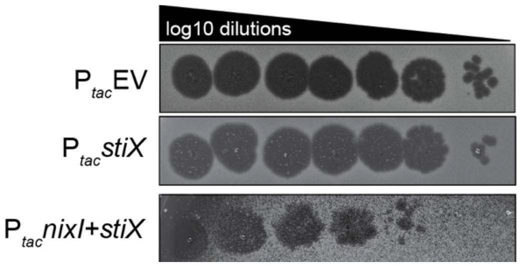


**Figure S8. Nix+StiX induction reduces ICP1 plaquing.** 10-fold dilutions of ICP1 (black) on a lawn of *V. cholerae* (grey) supplemented with inducer, expressing an empty vector, *stiX* or *nixI*+*stiX*. Expression of *nixI* alone is bactericidal at all inducer concentrations tested, so no lawn is able to form and thus was not included in this assay.

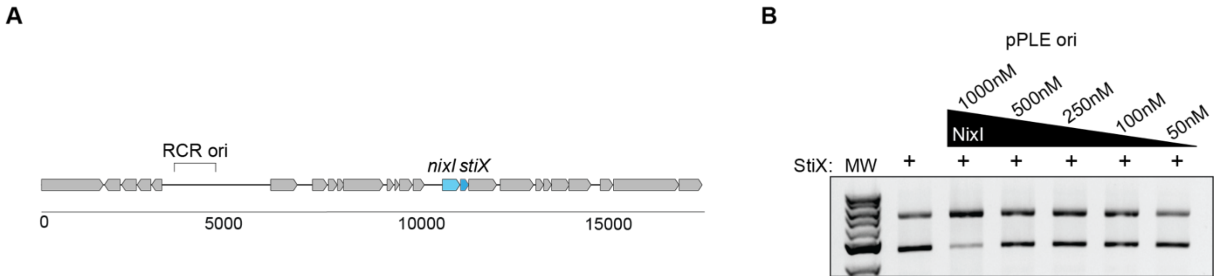


**Figure S9. NixI and StiX do not cleave PLE's rolling circle origin of replication, including PLE's putative packaging site** (A) Gene map of PLE 1 showing relative position of PLE's RCR origin (ori) (B) Cleavage assay with a plasmid substrate containing 500bp stretching across PLE's RCR origin, with 500nM StiX. MW is the molecular weight ladder. Plasmid incubated only with StiX is shown as a control.

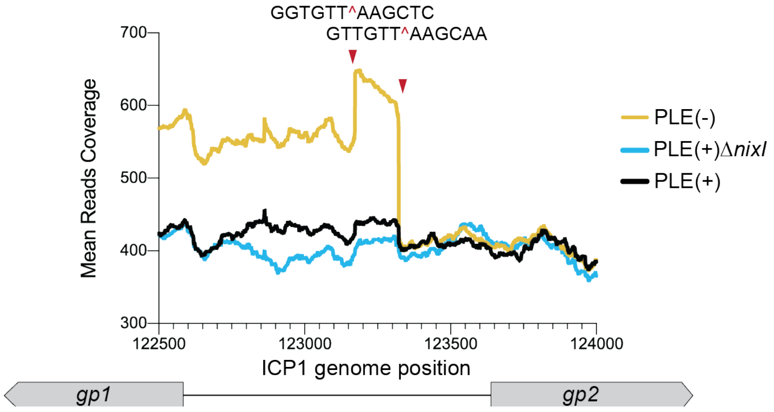


**Figure S10. PLE inhibits packaging of ICP1's genome independent of NixI**. Total reads as the average of three biological replicates across ICP1's putative *pac* site during infection of a PLE(-), PLE(+) and PLE(+)Δ*nixI* host, indicating that packaging is inhibited by PLE and not restored during infection of a Δ*nixI* host, suggesting an additional mechanism that inhibits packaging of ICP1's genome.

Supplementary Table S1: Strains used in this study

| Strain | Description* | Source |
|---|---|---|
| KDS6 | *V. cholerae* E7946 | Lab collection |
| KDS103 | *V. cholerae* E7946 containing PLE 1 $Kan^R$ | (1) |
| KDS36 | *V. cholerae* E7946 containing PLE 1 | (1) |
| KDS162 | *V. cholerae* E7946 PLE1 Δ*nixI*::frt, $Kan^R$ | (2) |
| KL228-229 | *V. cholerae* E7946 PLE1 Δ*nixI* pKL06: $P_{tac}$ Empty Vector, $Kan^R$, $Cm^R$ | This study |
| KL230-231 | *V. cholerae* E7946 PLE1 Δ*nixI* pKL06: $P_{tac}$*nixI*, $Kan^R$, $Cm^R$ | This study |
| KDS163 | *V. cholerae* E7946 PLE1 Δs*tiX*::frt, $Kan^R$ | This study |
| KDS196 | *V. cholerae E7946* pKL06 empty vector $Cm^R$ | (2) |
| KDS211 | *V. cholerae E7946* pKL06: $P_{tac}$*nixI* $Cm^R$ | This study |
| KL20 | *V. cholerae E7946* pKL06: $P_{tac}$*nixI* +*stiX* $Cm^R$ | This study |
| KL160-161 | *V. cholerae E7946* pKL06: $P_{tac}$*nixI* $^{N95A}$ $Cm^R$ | This study |
| KDS212 | *V. cholerae E7946* pKL06: $P_{tac}$*stiX* $Cm^R$ | This study |
| KL591-592 | *V. cholerae E7946* pKL06: $P_{tac}$*nixI* homolog from *V. cholerae* YB2_A06*,* $Cm^R$ | This study |
| KL593-594 | *V. cholerae E7946* pKL06: $P_{tac}$*nixI* homolog from *V. parahaemolyticus* s042*,* $Cm^R$ | Thus study |
| ICP1 | ICP1_2006_E ΔCRISPR ΔCas2-3 | (3) |
| KL249-250 | *V. cholerae* E7946 midiPLE, ΔlacZ::EV, pKL06 $P_{tac}$-*repA*, $Kan^R$, $Spec^R$, $Cm^R$ (RepA plasmid expression construct in strain with midiPLE and empty vector construct in lacZ locus) | This study |
| KL251-252 | *V. cholerae* E7946 midiPLE, ΔlacZ::$P_{tac}$-*nixI* , pKL06 $P_{tac}$-*repA*, $Kan^R$, $Spec^R$, $Cm^R$ (RepA plasmid expression construct in strain with midiPLE and inducible *nixI* in lacZ locus) | This study |
| KL253-254 | *V. cholerae* E7946 midiPLE, ΔlacZ:: $P_{tac}$-*nixI*+*stiX*, pKL06 $P_{tac}$-*repA*, $Kan^R$, $Spec^R$, $Cm^R$ (RepA plasmid expression construct in strain with midiPLE and inducible *nixI*+*stiX* in lacZ locus) | This study |
| KL133 | *E. coli* BL21 pLysS with IPTG inducible His-SUMO-*nixI* from PLE 1, $Kan^R$ for expression vector, $Cm^R$ for pLys vector (purification of NixI) | This study |
| KL124 | *E. coli* BL21 His-SUMO-*stiX*, $Kan^R$ (purification of StiX) | This study |
| KL171 | *E. coli* BL21 His-SUMO-*nixI*$^{N95A}$, $Kan^R$ (purification of catalytically inactive NixI) | This study |
| KL285 | *E. coli* pEV (pUC19 derivative plasmid) for empty vector control for nuclease assays, $Cm^R$ | This study |
| KL286 | *E. coli* pGp121 (pUC19 derivative plasmid), *gp121* region from ICP1 2006_E, $Cm^R$ | This study |
| KL287 | *E. coli* pPLE ori (pUC19 derivative plasmid), containing PLE 1's putative RCR ori and *pac* site, $Cm^R$ | This study |

| Strain | Description* | Source |
|---|---|---|
| KL645 | *E. coli* pGp121 (pUC19 derivative plasmid), *gp156* region from ICP1 2006_E, $Cm^R$ | This study |

* $Kan^R$ = Kanamycin resistance cassette, $Spec^R$ = Spectinomycin resistance cassette, $Cm^R$=Chloramphenicol resistance cassette

Supplementary Table S2: Primers used in this study

| Primer | Sequence | Purpose | Source |
|---|---|---|---|
| zac14 | AGGGTTTGAGTGCGATTACG | qPCR PLE | (1) |
| zac15 | TGAGGTTTTACCACCTTTTGC | qPCR PLE | (1) |
| zac68 | CTGAATCGCCCTACCCGTAC | qPCR ICP1 as well as gp62 EMSA probe | (1) |
| zac69 | GTGAACCAACCTTTGTCGCC | qPCR ICP1 as well as gp62 EMSA probe | (1) |
| KL119 | cgaacagattggaggtATGCCAACACAAAAGACGT | *nixI* with pSUMO overhangs | This study |
| KL120 | tggtggtggtgctcgaTTTAAATTTCTAGCCGATCT | *nixI* with pSUMO overhangs | This study |
| KL143 | gaacagattggaggtATGATAACCATCGAACTG | stiX with pSUMO overhangs | This study |
| KL144 | gtggtggtggtggtgctcgaTCATTCTTTTGTTTCCTTAAATTTG | stiX with pSUMO overhangs | This study |
| KL170 | CAAGAGgccTTAAAAGAAACTTCTAATTTTTACTTAG | *nixI* $^{N95A}$ point mutation | This study |
| KL171 | CTTTTAAggcCTCTTGGTGACTACACCAC | *nixI* $^{N95A}$ point mutation | This study |
| KL226 | CTATGACCATGATTACGCCAGTCGATGTTCTAAAGGGTATCG | Gp121 cut site overhangs to p958 | This study |
| KL227 | GTTGTAAAACGACGGCAGCCTCTGATAGTGCTTCGT | Gp121 cut site overhangs to p958 | This study |
| KL233 | CTATGACCATGATTACGCCAAAGTTACCTCAGTGTCGTTAG | Gp156 cut site overhangs to p958 | This study |
| KL234 | GTTGTAAAACGACGGCCTCATCCATCCACACAAACCT | Gp156 cut site overhangs to p958 | This study |
| KL224 | GACGTTGTAAAACGACGGCCTATATTTGAGTATAAGTGTGGTGC | Clone PLE's putative pac site onto vector p958 | This study |

| Primer | Sequence | Purpose | Source |
|---|---|---|---|
| KL225 | GCTATGACCATGATTACGCCACTGGAAA TAATGACAATTGAGATTGC | Clone PLE's putative pac site onto vector p958 | This study |
| KL241 | GATGGAAGGGCTAACGGT | 200bp EMSA probe of gp121 | This study |
| KL242 | CATCTCCGTGATTACCTTGGG | 200bp EMSA probe of gp121 | This study |
| KS951 | CTGCTAAGGAGGTAACAACAAGATGCCA ACACAAAAGACGTTAGACC | *nixI* pKL06 overhangs | This study |
| KS962 | ctctcatccgccaaaacagcTTAAATTT CTAGCCGATCTATTTTAAATCG | *nixI* pKL06 overhangs | This study |
| KS966 | ctctcatccgccaaaacagcTCATTCTT TTGTTTCCTTAAATTTGTTGTTG | R oligo stiX pKL06 overhangs | This study |
